# Integrating computational and experimental biophysics reveals novel insights into the RAD51-BRC4 interaction

**DOI:** 10.1101/2025.03.07.642044

**Authors:** Veronica Bresciani, Francesco Rinaldi, Pedro Franco, Stefania Girotto, Andrea Cavalli, Julian D. Langer, Matteo Masetti, Mattia Bernetti

## Abstract

GRAPHICAL ABSTRACT

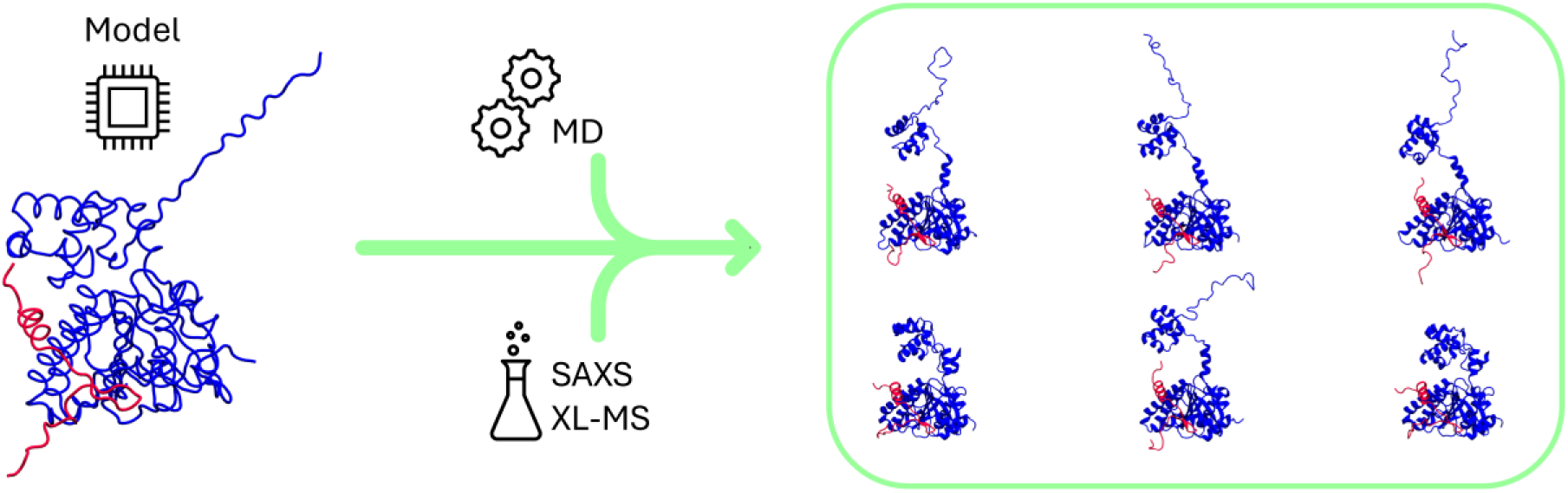

The interaction between the RAD51 and BRCA2 proteins is central for homologous recombination, a crucial pathway ensuring high-fidelity DNA repair. Recruitment of RAD51 involves eight highly conserved regions on BRCA2, named BRC repeats. To date, only the interaction between the fourth BRC repeat (BRC4) and the RAD51 C-terminal domain has been structurally characterized, while the complex of full-length RAD51 with the peptide still remains elusive. Here, we report an integrative experimental and in silico approach to reconstruct the conformational ensemble in solution for full length RAD51 in complex with BRC4. We combined AlphaFold2, crosslinking mass spectrometry (XL-MS) and small angle x-ray scattering (SAXS) data with molecular dynamics simulations (MD). These data show that the full-length RAD51-BRC4 complex is a mixture of compact and elongated conformations. Detailed analysis of the reweighted ensemble, achieved through the maximum entropy principle, identifies key residues at the N-terminal-BRC4 interface mediating complex conformational dynamics. Our evidence provides robust atomic-level insights into the interaction of RAD51 and BRC4. These findings are crucial for understanding the molecular features underlying the recognition between RAD51 and BRCA2, which are essential for developing therapeutic intervention strategies in cancer treatment.

## INTRODUCTION

Homologous recombination (HR) is an essential process of Synthesis/Growth 2 (S/G2) phases, ensuring the high fidelity fixing of very deleterious DNA lesions known as double strand breaks (DSBs) (1). HR requires a series of well-coordinated steps involving multiple proteins (2). Among these, BRCA2 and RAD51 play a crucial role (3). RAD51 is a recombinase which displays the tendency to assemble into fibril-like oligomers which are fundamental to enable the strand exchange activity, an essential step of HR (4–6). BRCA2 has been demonstrated to interact with numerous factors involved in HR, likely functioning as hub to efficiently orchestrate this critical DNA repair process (3, 7, 8). Although the three-dimensional structure of BRCA2 has not been solved yet, biochemical evidence proved that it can interact with RAD51 through eight highly conserved sequences, referred to as BRC repeats (3, 9, 10). Very limited structural information is available on this protein-protein interaction, however BRC repeats are supposed to act synergistically allowing the recruitment of RAD51 in the cytosol and directing it to damaged DNA sites (9, 11). X-ray crystallography structural studies have already allowed to elucidate the interaction between the RAD51 C-terminal (C-ter) domain and the fourth BRC repeat (BRC4) which was reported to harbor the highest affinity for RAD51 (11). This work unveiled the FXXA and LFDE domains as crucial for the binding of BRC4 to RAD51 (11, 12). Nevertheless, the complex of full-length RAD51 (FL-RAD51) and BRC4 remains elusive to conventional structure determination techniques, thus hindering the ability to uncover potential interactions between BRC4 with the RAD51 N-terminal domain (N-ter), as well as the structural changes that BRC repeats binding might induce in this region promoting RAD51 filaments depolymerization (3, 13, 14). This information would be crucial to provide novel insights useful to understand how mutations within BRC4 can promote cancer onset as well as novel potential druggable pockets to develop innovative disruptors of the BRCA2-RAD51 interaction to leverage the synthetic lethality paradigm in combination with PARP inhibitors (PARP-i) (15).

The first attempt to shed light on the FL-RAD51-BRC4 complex dates to 2013 (16). In this work, Subramanyam and co-workers were able to reconstruct the complex by coupling homology modeling, molecular dynamics simulations and biophysical/biochemical evidence, thus suggesting a series of hydrogen bonds subsisting between BRC4 and the RAD51 N-ter (16). More recently, modelling of AlphaFold2 derived structures based on Small Angle X-Ray Scattering (SAXS) studies on the RAD51-BRC4 complex suggested that in the presence of BRC4 the RAD51 N-ter, displays complex conformational dynamics possibly behaving as an intrinsically disordered region without accounting the possible BRC4-RAD51 N-Ter domain interactions (13). This observation provided a possible explanation of the difficulties encountered in obtaining the X-ray crystal structure of the RAD51-BRC4 complex (13). Indeed, achieving high resolution structural insights into biomolecules behaving as flexible systems (e.g. intrinsically disordered proteins (IDPs)) can be particularly challenging (13). Moreover, it should be accounted that such dynamic entities can adopt multiple conformations in solution which can be better described as structural ensembles (13). In this sense, SAXS experiments can provide valuable low-resolution insights of biomolecules directly in solution, typically in terms of generic properties such as system size and degree of structural compactness which are particularly useful to describe ensembles of IDPs or nucleic acids (e.g. RNA) (17, 18).] Specifically, this experimental information can be combined with Molecular Dynamics simulations (MD) to study the conformational dynamics of biologically relevant systems with an atomic-level description under near-physiological conditions (19–22). However, drawbacks in the underlying physical models (i.e. the force fields), and sampling limitations hinder the predictive capacities of this simulative approach (23). On the one hand, the latter aspect can be mitigated using approaches specifically devised to improve the sampling of the configurational space, namely enhanced sampling methods (24–26). On the other hand, addressing the aspect of force field inaccuracies, which affect results with systematic errors, is a highly challenging task (23, 27), especially for biomolecular systems displaying particularly complex structural dynamics, such as IDPs or RNA (28–31). In this regard, the combination of simulations and experiments can be beneficial in overcoming the respective limitations, allowing an accurate reconstruction of atomic-scale conformational ensembles in solution (20–22, 32, 33). Moreover, other orthogonal experimental approaches can be further integrated in MD simulations to refine the structural dynamics of the system in solution (20). For instance, crosslinking mass spectrometry (XL-MS) can pinpoint residues located within specific distances in the three-dimensional structure of proteins, thus allowing to probe protein-protein interactions and providing both a validation tool for AI generated structure models and valuable restraints in MD simulations (34, 35).

In this work, we exploit state-of-the-art computational approaches and experimental data derived from SAXS and XL-MS methods to reconstruct the conformational ensemble of the RAD51 protein in complex with BRC4, in solution (**Figure 1**). Our protocol leverages artificial intelligence (AI) methods, namely AlphaFold2, to generate an initial guess of the RAD51-BRC4 complex. The conformational dynamics of this structure is then explored via enhanced-sampling MD approaches, and the structural ensemble of the RAD51-BRC4 complex in solution is finally reconstructed by integrating SAXS and XL-MS experimental data in this pipeline (**Figure 1**). Through our procedure, we first show how accounting for the solvent contribution in the calculation of SAXS spectra from MD-sampled structures is necessary to drive the system towards a single-structure model compatible with the SAXS experimental spectrum. Then, we integrate information from XL-MS data into enhanced sampling simulations to effectively explore the system’s conformational ensemble while optimizing simulation efficiency. The resulting ensemble is finally reweighted to improve its agreement with SAXS data. Such a procedure revealed that a mixture of both compact and more elongated conformations of the system is required to accommodate experimental data. A finer characterization of the reconstructed ensemble allows us to determine residues that are critical in governing the interaction between the interfaces of the BRC4 repeat and RAD51’s N-ter.

**Figure 1.**
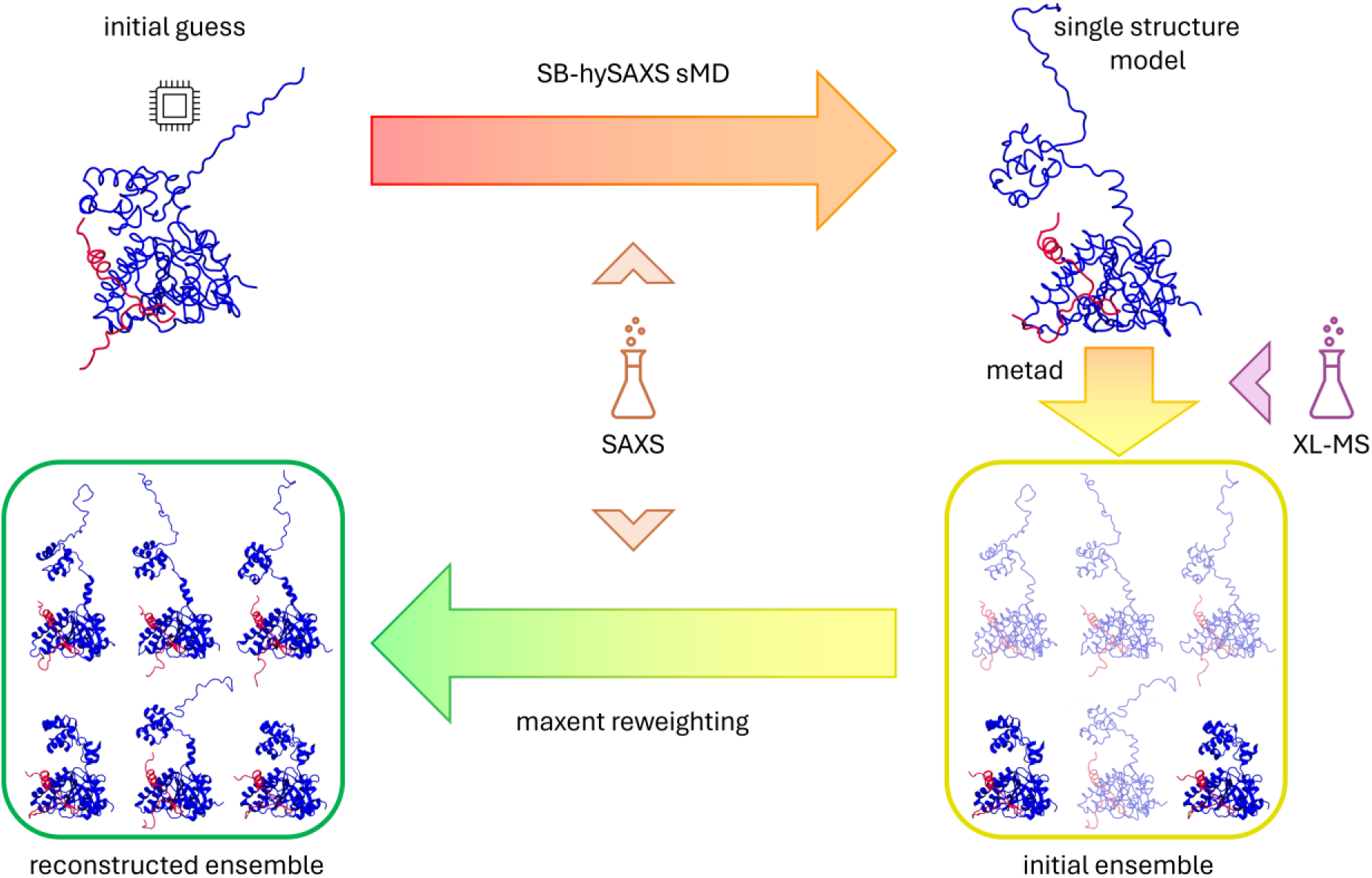
Schematic depiction of the pipeline presented in this work. An initial structural guess for the full RAD51-BRC4 complex is generated through the structure prediction tool AlphaFold2. Steered MD simulations (red to orange arrow) and then used to increase compatibility with reference SAXS experimental data (orange flask). Metad simulations (orange to yellow arrow) started from the improved structure are used to generate an ensemble of diverse configurations of the RAD51-BRC4 complex (yellow rounded rectangle, different shading denotes different weights). Newly generated XL-MS data (purple flask) are integrated at this stage to optimize sampling. Finally, the generated ensemble (prior, yellow rounded rectangle) is reweighted through the maximum entropy principle (yellow to green arrow) to identify an ensemble of structures (posterior, green rounded rectangle) in agreement with experimental SAXS data.

## MATERIAL AND METHODS

### His-RAD51 [F86E, A89E] expression and purification and peptide synthesis

Monomeric His-RAD51 [F86E, A89E] was expressed and purified as previously described in (13).

BRC4 peptide (NH2-KEPTLLGFHTASGKKVKIAKESLDKVKNLFDEKEQ-COOH) was synthesized by Thermofisher Scientific, while Biotinylated BRC4 (BioBRC4) (Bio-Ahx-KEPTLLGFHTASGKKVKIAKESLDKVKNLFDEKEQ-COOH) was synthesized by Peptide Protein Research Ltd.

### AlphaFold predictions

All predictions reported in this work were generated through AlphaFold 2.3.2 run as a Singularity container. The model preset used for prediction was set to multimer, enabling Amber relaxation, which resolves remaining structural violations and clashes in the predicted structure through an iterative restrained energy minimization via gradient descent with the Amber ff99SB force field (36), only for the highest ranked model. Database preset was set to full database (full_dbs databases: bfd, mgnify (mgy_clusters_2022_05), pdb_mmcif, pdb_seqres, pdb70, UniRef30_2023_02, uniprot, uniref90). Graphs of multiple sequence alignment (MSA) coverage and sequence similarity predicted local distance difference test (pLDDT) and predicted aligned error (PAE) were generated through Python scripts adapted from https://raw.githubusercontent.com/jasperzuallaert/VIBFold/main/visualize_alphafold_resultr.p y and https://raw.githubusercontent.com/busrasavas/AFanalysis/main/AFanalysis.py

### Static Light Scattering

Static light scattering (SLS) analyses were performed on a Viscotek GPCmax/TDA (Malvern, UK) instrument, connected in tandem with a series of two TSKgel G3000PWxl size-exclusion chromatography columns (Tosoh Bioscience) as already described in (13). For all the experiments the system was equilibrated with buffer containing 20 mM Hepes pH 8.00, 100 mM Na_2_SO_4_, 5% glycerol. Monomeric RAD51 [F86E, A89E] at 24 µM (0.94 mg/mL) was incubated for 1 hour on a thermo-block at 25 °C, in the absence (buffer only) or presence of BRC4 or biotinylated BRC4 peptides (respectively dissolved in the running buffer (stock concentration 1 mM) and in 100% DMSO (stock concentration 2 mM)) in a 4-fold higher stoichiometric excess (final DMSO concentration for BioBRC4 = 5%). Data analysis was performed using Viscotek software, calibrating the instrument with Bovine Serum Albumin (BSA) at 5 mg/mL. Data were exported as .csv files and re-graphed using GraphPad Prism 10.

### Biolayer Interferometry

Biolayer interferometry (BLI) experiments were performed utilizing an Octet K2 system (Sartorius). All BLI experiments were carried out in an assay buffer containing 20 mM Hepes pH 8.00, 100 mM Na_2_SO_4_, 5% glycerol, 0.05% Tween 20, 0.1% PEG8000 and 0.5 mM Sodium Deoxycholate. The following protocol was applied: 60 sec baseline, 240 sec loading, 240 sec baseline, 180 sec association, 180 sec dissociation. For every step, shaking at 1000 rpm was enabled and temperature set at 25°C. Streptavidin Octet biosensors (18-5019, Sartorius) were initially dipped into assay buffer to record an initial baseline of 60 seconds. BioBRC4 was solubilized in 100% DMSO at a 2 mM concentration, diluted at a final 1 µM concentration in assay buffer and then immobilized to the Streptavidin sensor through 240 seconds loading step. After the loading stage, a second baseline of 240 seconds was recorded in wells containing only assay buffer to verify the stability of signal and remove unbound peptide. For each experiment, sensors were subsequently dipped for 180 seconds into wells containing HisRAD51 [F86E, A89E] (460 nM, 230 nM, 115 nM) to measure association signals and finally moved to wells containing only assay buffer to assess complex dissociation. Two replicates were run for each His-RAD51 [F86E, A89E] concentration. BLI experiments were carried out including a double reference: a reference well (where only immobilized bioBRC4 was present on the sensor and no analyte (0 nM) during association) and reference sensor (where no bioBRC4 was immobilized on the sensor and HisRAD51 [F86E, A89E] concentration was matched during association). Data were analyzed using Octet Analysis Studio 12.2 subtracting to recorded sensorgrams the signals of both the reference well and reference sensors to remove signals due to aspecific binding. Recorded sensorgrams were corrected by aligning to the average of the second baseline steps, applying Savitzky–Golay filtering and inter-step correction based on the second baseline step. To calculate R_max_, the response at the end of the association phase (170-175 sec) was extrapolated. All data presented data were exported to .csv files and re-graphed using GraphPad Prism 10.

### Cross-linking: sample preparation and reaction conditions set up

For crosslinking with bis(sulfosuccinimidyl)suberate (BS3), purified His-RAD51[F86E, A89E] concentration was adjusted to 24 µM (=0.98 mg/mL) and mixed with an equimolar concentration of biotinylated BRC4 peptide (previously solubilized as a 2 mM stock in 100% DMSO) with a final DMSO concentration lower than 5% (v/v). The mixture was incubated for 30 minutes at 20°C on a thermoblock to allow complex formation and then crosslinked with a final concentration of 1 mM BS3 (previously solubilized in PBS as a 50 mM stock). After 30 minutes at the same temperature, 4x Laemmli Sample Buffer (BioRad #1610747) was added to cross-linked samples and boiled at 95°C for 5 minutes. For crosslinking with 1-Ethyl-3-(3-Dimethylaminopropyl) carbodiimide, Hydrochloride (EDAC) purified His-RAD51[F86E, A89E] at the same concentration reported above (24 µM (=0.98 mg/mL)) was mixed with a 2-fold stoichiometric excess of biotinylated BRC4 peptide (48 µM, final DMSO concentration lower than 5%). The mixture was incubated for 1 hour at 20°C to allow complex formation and then crosslinked with a final concentration of 0.2% (w/v) EDAC (previously solubilized in DMSO as a 10% w/v stock) for 1 hour at the same temperature. 4x Laemmli Sample Buffer (BioRad #1610747) was then added to cross-linked samples and boiled at 95°C for 5 minutes.

### Coomassie Blue and Western Blot analyses on cross-linked samples

Efficiency of crosslinking reactions was evaluated through Coomassie blue staining and western blot analysis. Prepared cross-linked samples were resolved using an 4-15% SDS-PAGE gel (Criterion™ TGX™ Precast Midi Protein Gel), which was stained using Page Blue Protein Staining solution (Thermo Fisher, 24620) according to manufacturer protocols. Images were acquired using a gel Doc EZ Imager (Biorad). For Western Blot (WB), an amount equivalent to 50 ng of His-RAD51[F86E, A89E], and 5 ng of bioBRC4 was loaded. Sample was run on a 12% SDS-PAGE gel (Criterion™ TGX™ Precast Midi Protein Gel) and then electrophoretically transferred to TransBlot Turbo nitrocellulose membranes (Midi size, Biorad) using a Transblot Turbo apparatus (Biorad, set at 25V, 1.0A, 30 minutes). Membranes were blocked for 1 hour at room temperature in 5% Milk TBS-T and after one wash with TBS-T were incubated for 1 hour at room temperature with Streptavidin HRP to detect the biotin moiety of bioBRC4 (1:3000 dilution in 3% BSA-TBS-T). After three washes in TBS-T, chemiluminescence was detected using the Clarity Western ECL substrate (Biorad, #1705061), and images were recorded using a ChemiDoc MP Imaging System (Biorad).

### LC-MS analysis

Crosslinking samples in Laemmli Sample Buffer were digested following the S-Trap protocol (37) (Protifi) with minor adaptations and purified using solid-phase extraction. Briefly, a volume of sample containing 100 µg of proteins was diluted to a final volume of 100 µl with 50 mM ammonium bicarbonate (ABC) and proteins were reduced by addition of 4 µl TCEP for 30 min at 37°C with mild agitation. Cysteine alkylation was performed with 8 µl iodoacetamide (IAA) for 30 min in the dark with mild agitation. The samples were then acidified with 12 µl 12% phosphoric acid, diluted with 750 µl of the S-Trap binding buffer (1 M triethylammonium bicarbonate (TEAB) Buffer and Methanol (10:90)) and transferred to S-Trap mini columns (Protifi). SDS removal was conducted by three washing cycles with 400 µl S-Trap binding buffer prior to digestion. MS-grade Trypsin (Serva) was added in an enzyme to protein ratio 1:50 using 125 µl digestion buffer (50mM ABC, pH 8,5) as media and the column incubated overnight at 37°C with light agitation. Peptides were eluted stepwise in 80 µl 50mM ABC, 0,2% formic acid, 50% acetonitrile (ACN). The pooled fractions were diluted 1:1 with 0,1% trifluoroacetic acid (TFA) and desalted with C18-SPE cartridges (Biotage). After equilibration with 2 ml ACN, 1 ml 50% ACN/1% acetic acid and 2 ml 0,1% TFA the samples were loaded onto the cartridge, washed with 2 ml 0,1% TFA and eluted with 1 mL 80% ACN/0,1% TFA. The eluted fractions were dried using an Eppendorf concentrator (Eppendorf) and stored at −20°C before analysis. Dried peptides were reconstituted in 5% ACN with 0.1% formic acid (FA). Peptides were loaded onto an Acclaim PepMap C18 capillary trapping column (particle size 3 µm, L = 20 mm) and separated on a ReproSil C18-PepSep analytical column (particle size = 1,9 µm, ID = 75 µm, L = 25 cm, Bruker Coorporation, Billerica, USA) using a nano-HPLC (Dionex U3000 RSLCnano) at a temperature of 55°C. Trapping was carried out for 6 min with a flow rate of 6 μL/min using a loading buffer composed of 0.05% trifluoroacetic acid in H_2_O. Peptides were separated by a gradient of water (buffer A: 100% H_2_O and 0.1% FA) and acetonitrile (buffer B: 80% ACN, 20% H_2_O, and 0.1% FA) with a constant flow rate of 400 nL/min. The gradient went from 4% to 48% buffer B in 45 min. All solvents were LC-MS grade and purchased from Riedel-de Häen/Honeywell (Seelze, Germany). Eluting peptides were analyzed in data-dependent acquisition mode on an Orbitrap Eclipse mass spectrometer (Thermo Fisher Scientific) coupled to the nano-HPLC by a Nano Flex ESI source. MS1 survey scans were acquired over a scan range of 350–1400 mass-to-charge ratio (m/z) in the Orbitrap detector (resolution = 120k, automatic gain control (AGC) = 2e5, and maximum injection time: 50 ms). Sequence information was acquired by a “ddMS2 OT HCD” MS2 method with a fixed cycle time of 5 s for MS/MS scans. MS2 scans were generated from the most abundant precursors with a minimum intensity of 5e3 and charge states from two to eight. Selected precursors were isolated in the quadrupole using a 1.4 Da window and fragmented using higher-energy collisional dissociation (HCD) at 30% normalized collision energy. For Orbitrap MS2, an AGC of 5e4 and a maximum injection time of 54 ms were used (resolution = 30k). Dynamic exclusion was set to 30 s with a mass tolerance of 10 parts per million (ppm). Each sample was measured in duplicate LC-MS/MS runs. MS raw data were processed using the MaxQuant software (38) (v2.6.5.0) with customized parameters for the Andromeda search engine. Spectra were matched to a FASTA file containing the BRCA2 and RAD51 sequences downloaded from UniProtKB (October 2023), a contaminant and decoy database. The RAD51 sequence was modified to include the His-Tag on the N-terminus (MGSSHHHHHHSSGLVPRGSHMLEDP-). A minimum tryptic peptide length of seven amino acids and a maximum of two missed cleavage sites were set. The crosslinker BS3 was chosen from the default list of available crosslinkers. Precursor mass tolerance was set to 4.5 ppm and fragment ion tolerance to 20 ppm, with a static modification (carbamidomethylation) for cysteine residues. Acetylation on the protein N-terminus and oxidation of methionine residues were included as variable modifications. A false discovery rate (FDR) below 1% was applied at crosslink, peptide, and modification levels. Search results were imported to xiVIEW (39) for subsequent analysis. After stringent manual curation of the spectra, only inter-protein crosslinks with spectral scores above 30 were selected for fitting on the structures. Crosslinks were fitted on the His-Tag-RAD51 and BRC4 and on the untagged RAD51 and BRC4 structures, generated through Alphafold 2.3, using xiVIEW.

### SAXS spectra calculation

Experimental observables can be predicted from structures through forward models, i.e. equations relating measured quantities with structural features. Herein, the calculation of SAXS spectra from both AmphaFold2 and MD-sampled structures was performed with the PLUMED library (40, 41), version 2.9, via the Integrative Structural and Dynamical Biology (ISDB) module (42). In particular, the SAXS spectrum of a given three-dimensional structure can be calculated according to the following equation:

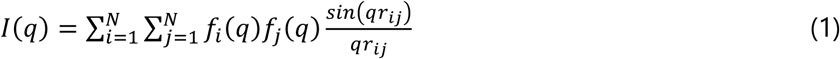

where the scattered intensity *I* at a certain value of the momentum transfer *q* is obtained from the pairwise distance *r*_ij_ between all pairs of the *N* atoms in the biomolecules and the corresponding atomic scattering factors *f*_*i*_(q) and *f*_*j*_(q).

Calculating the pairwise distance between each pair of atoms in a macromolecular system at atomistic resolution is computationally demanding. This becomes particularly critical if the calculation has to be repeated many times, e.g. on a large structural ensemble or on-the-fly during MD simulations. For this reason, a coarse-grained representation can improve the efficiency of the calculation. Specifically, the atomistic representation can be simplified by grouping together a given number of atoms into pseudoatoms, also called beads. This results in a system representation comprising *M* beads, with typically *M* ≪ *N* and the computed scattering intensity can be expressed as:

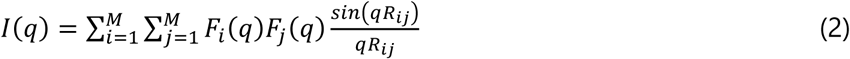

where *R*_*ij*_is the distance between pairs of beads. Notably, this implies that scattering factors *F* associated with the pseudoatoms are available, i.e. have been purposely parameterized. Most interestingly, a great advantage of this strategy is that it can be exploited in a hybrid coarse-grain/all-atom fashion. Specifically, the MD simulations can be conducted at a fully atomistic resolution, while resorting to a coarse-grained representation for the sole purpose of computing efficiently the SAXS spectra (hySAXS scheme).

The MARTINI force field is one of the most popular coarse-grained models for biolomolecules (43). In such a paradigm, amino acid residues are mapped into a varying number of beads, ranging from 1 (e.g. alanine) to 5 (e.g. tryptophane). Scattering factors of MARTINI beads have been parameterized for both protein and nucleic acid systems, allowing for hySAXS scheme simulations exploiting MARTINI as the coarse-grained model (hereafter referred to as MT-hySAXS). Recently, an alternative coarse-grained model for SAXS spectra calculation was introduced, named Single-Bead (44, 45). In the protein context, this representation replaces each amino acid with a single bead, hence the name. Despite being more simplified than the MARTINI representation, thus further improving the efficiency of SAXS spectra calculation, this model retains comparable accuracy, particularly for *q* values lower than 0.3 A^2^ (45). Remarkably, the Single-Bead approach (SB-hySAXS) features a solvent layer contribution term (45) in the definition of scattering factor *F* that allows accounting for the effect of solvation on solvent-exposed atoms in the biomolecule.

### Steered MD simulations

All MD simulations were carried out at the atomistic level using the GROMACS MD engine (46), using the Amber ff19SB force field (47) for the proteins, and the OPC model for water molecules (48). The full structure of the RAD51-BRC4 complex predicted with AlphaFold2 was inserted in a dodecahedron-shaped box, with edges distant 15 Å from the biomolecules. The box was then filled with OPC waters and the system was neutralized and brought to physiological ionic concentration (0.15 M) with NaCl using Joung and Cheatham ions (49). The system was energy minimized using the steepest descent method and subsequently equilibrated via a 1.2 ns simulation in the NVT ensemble with position restraints of 239 kcal/mol on protein heavy atoms (of which, 400 ps were conducted at 100 K, 400 ps at 200 K, and the last 400 ps at 300 K) and 800 ps in the NPT ensemble (of which, 400 ps were conducted with protein heavy atoms restraints, while the last 400 ps only featured alpha carbon atoms restraints) using the V-rescale thermostat (50) with time constant 0.1 for temperature control and C-rescale barostat (51) with time constant of 0.1 for pressure control.

Steered MD (sMD) simulations were carried out via the MOVINGRESTRAINTS directive in PLUMED 2.9 (40, 41). Three replicates of 15 ns each were performed using the Martini coarse grain representation for SAXS spectra calculations, and three replicates of 15 ns using the single-bead model.

A set of 19 SAXS intensities computed at different values of the momentum transfer *q*, in the range 0.00-0.30 Å^-1^, was used as collective variable (CV). During the first 10 ns of sMD, the positions of the harmonic restraints were linearly interpolated from the values of the SAXS intensities computed on the initial structure, i.e. the AlphaFold2 model, to the experimental ones. The SAXS experimental spectrum for the RAD51-BRC4 complex was taken from the Small Angle Scattering Biological Data Bank (SASBDB), and deposited with code SASDQT9. To reduce the influence of experimental noise, the experimental values were taken after performing a 51-point running average on the experimental SAXS spectrum. The force constant was kept constant at a value of 10^6^ kJ/(mol*a.u.^2^) for the entire simulation.

### Metadynamics simulation

A 100 ns Well-Tempered Metadynamics (52) run in the NVT ensemble was carried out via PLUMED 2.9, using the radius of gyration as CV. Gaussians of width 0.05 Å and height 0.500 kcal/mol were deposited every 500 steps, using a bias factor of 10. The distances between Cα atoms of the four pairs of cross-linked lysine residues, as indicated by the XL-MS experiments, were restrained through a flat-bottom restraining potential, as implemented in the UPPER_WALLS PLUMED directive, positioned at a value of 30 Å (53) with a force constant of 23.90 kcal/(mol*Å^2^). Specifically, the restrained distances were the ones between RAD51’s K59 and BRC4’s K24, RAD51’s K59 and BRC4’s K29, RAD51’s K65 and BRC4’s K29, RAD51’s K71 and BRC4’s K29. Metad weights were computed a posteriori using the final bias.

### Maximum Entropy reweighting

The Maximum Entropy principle can be used to reweight ensembles (21, 54, 55) ensuring consistency with experimental data while inducing the least possible modification to the original distribution (the prior). In our case, the pool of conformations sampled by metadynamics was reweighted to bring the computed average SAXS spectrum into agreement with available experimental data.

Through this procedure, new weights *w*_t_ are assigned to each configuration *x*_t_ from the original ensemble comprising *N*_s_ structures (i.e. the frames in the MD trajectory), according to:

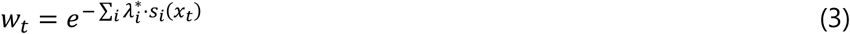

In this expression, *s*_i_(x_t_) is the value of the i^th^ observable (here the i^th^ intensity of a SAXS spectrum) computed for structure *x*_t_, and λ^*^_*i*_ is the corresponding Lagrangian multiplier that minimizes the Lagrangian function:

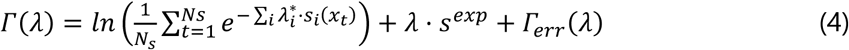

This minimization is the equivalent of Shannon’s Entropy maximization (56). As a result, the (weighted) average of the computed observable *s* along the reweighted trajectory is constrained to the measured experimental value *s^exp^*.

The last term in the equation is introduced to model experimental error, acting as a regularization term to reduce over-fitting, and can be defined as follows:

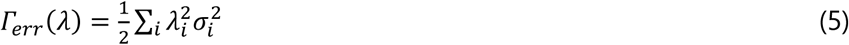

where a different *σ*_i_ value can be specified for each observable *s*_i_.

Herein, we used a prior ensemble generated via metad, characterized by uneven weights associated with the metad bias. In this case, the above formulations can be rewritten as:

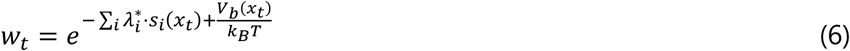

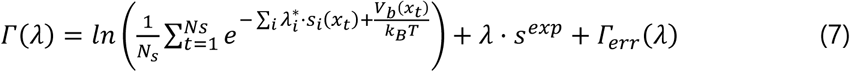

where *V*_b_(*x*t) is the metad bias at the end of the simulation recomputed for the coordinates of the *x*_t_^th^ frame, and *k*_B_*T* is the thermal energy. Herein, minimization was conducted through the optimize module in SciPy (57) using the BFGS (Broyden–Fletcher–Goldfarb–Shanno) method (58–60), and for simplicity we applied the same *σ* value to all observables, i.e. to all points in the spectrum.

Typically, only a fraction of the structures from the prior ensemble contribute effectively to the reweighted ensemble. The number of such structures, that carry significant weight, can be approximately estimated by calculating the Kish effective sample size:

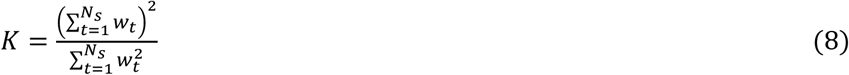

A parameter that is typically employed to assess overfitting in the context of SAXS spectra is the reduced χ^2^:

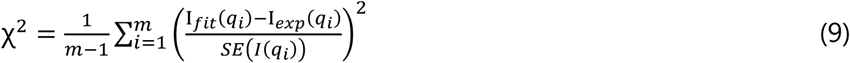

where I*_fit_*(*q_i_*) and I_exp_(*q_i_*) are the predicted (i.e. from the reweighted ensemble) and experimental intensities, and *SE*(*I*(*q_i_*)) the standard error on the intensities, at each *q_i_* value of the SAXS spectrum. χ^2^ values close to 0 indicate overfitting, while values around 1 indicate optimal fitting of the experimental data. Here, *SE*(*I*(*q_i_*)) included both the experimental and predicted uncertainties after applying error propagation.

### Trajectory analysis

The trajectory obtained from the Metadynamics simulation was analyzed through principal component analysis (PCA) carried out on the Cartesian coordinates of the system’s α carbons after aligning the snapshots on the α carbons of the C-ter domain. The first 5 PCs, explaining a cumulative variance of 93%, were used to compute a distance matrix on which we performed a weighted cluster analysis using the Quality Threshold (QT) algorithm, with a cutoff of 3 and the maxent weights. The cluster analysis, performed using the bussilab python package, was done separately on compact and extended structures, using an Rg value of 2.47 Å to discriminate between the two groups. All reported statistical errors were computed through standard bootstrapping with 400 iterations, after dividing the snapshots from the metad-generated trajectory in 10 blocks.

The ensemble of conformations was summarized and visualized by constructing a conformational space network, where the representative structure of each cluster served as a node. Edges in the network were assigned using a distance cutoff of 5, whereas the size of the nodes is proportional to the maxent weights. To better visualize the most populated clusters in the network, the weights were rescaled using a logistic function. The Pyvis-0.1.3.1 software (61) was used to represent the network, employing the BarnesHut (62) layout algorithm.

The contact analysis on the reweighted ensemble was performed using the compute_contacts module in MDTraj (63), using the closest-heavy scheme to compute minimum distances between pairs of residues. To define a contact, we employed a distance cutoff of 5 Å. The weighted average of the number of contacts was computed using the weights of the maxent-reweighted ensemble.

## RESULTS

### Probing the interaction between the RAD51’s N-ter and BRC4 combining AlphaFold2 and XL-MS

The crystal structure of the RAD51-BRC4 complex (PDB-ID: 1N0W) features RAD51’s C-ter domain but lacks information regarding RAD51’s N-ter domain, which was removed and replaced by a polypeptide linker connected to BRC4 (11). Superposition of this incomplete complex with the full-length *apo* structure of RAD51 (PDB ID: 5NWL), in the absence of DNA, clearly shows that the bound BRC4 repeat would clash with RAD51’s N-ter (**Figure 2A**), thus suggesting a conformational rearrangement of the RAD51 N-terminal domain when the BRC4 peptide binds (4, 11, 13).

**Figure 2.**
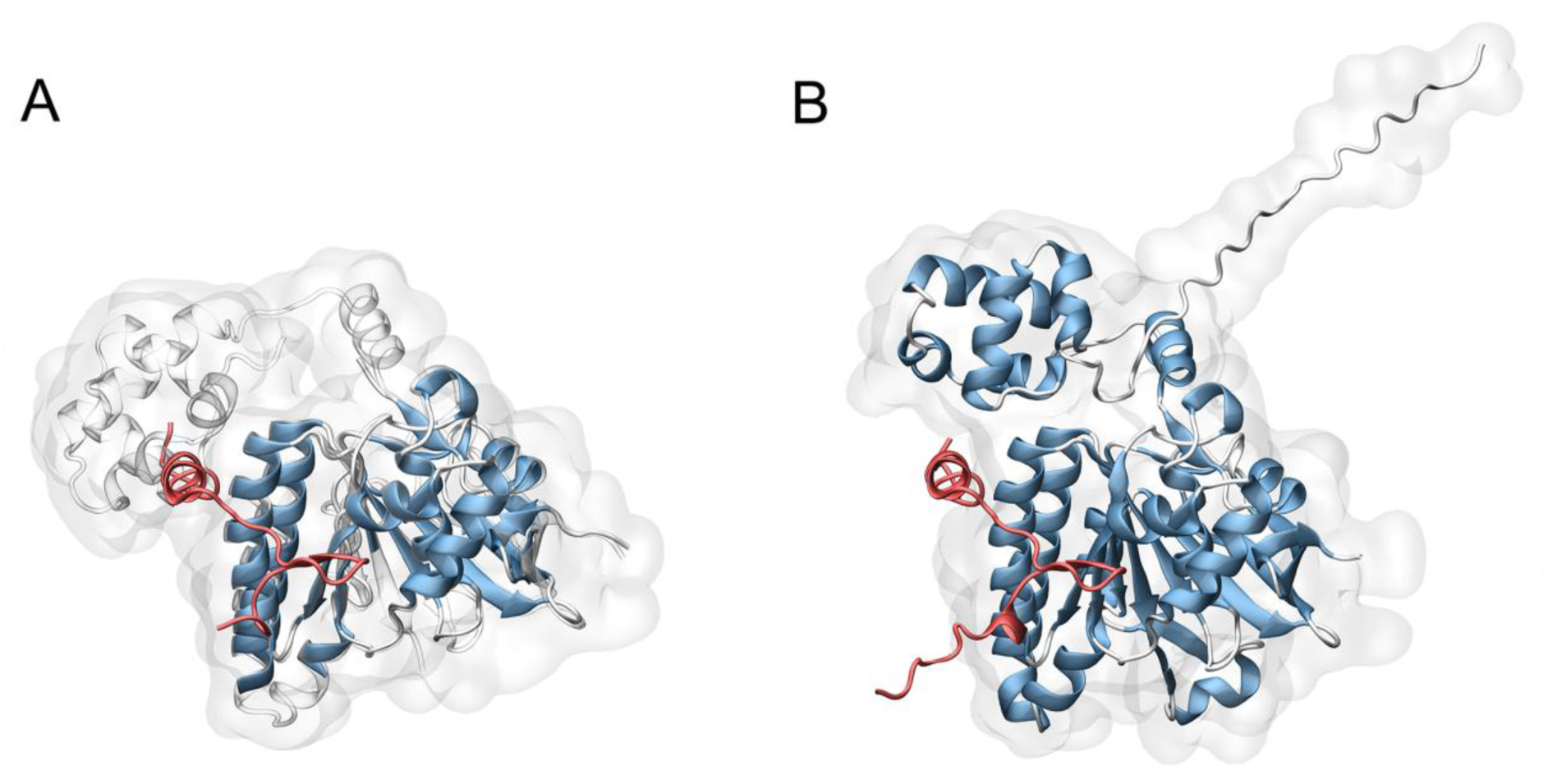
Structure of the RAD51-BRC4 complex. A) PDB entry 1N0W, featuring RAD51’s C-ter (in blue) and BRC4 (in red), superposed with PDB entry 5NWL, featuring RAD51’s C-ter and N-ter (in white) and lacking the BRC4 repeat. The superposition highlights how the N-ter domain and BRC4 are in overlap. B) The generated AlphaFold2 model, featuring RAD51’s C-ter and N-ter (in blue) and the BRC4 repeat (in red). In the prediction, the N-ter is shifted to preserve BRC4 binding and avoid overlap with the repeat.

The recent advent of AI algorithms to predict protein and protein complexes, such as AlphaFold2, has provided an unprecedented tool to generate initial guesses for structural investigations (36, 64, 65). Therefore, we used AlphaFold2 to generate a model of the FL-RAD51-BRC4 complex (**Figure 1**) (36). An important feature of AlphaFold2 is the implementation of scoring functions which enable the interpretation of the generated predictions as predicted local difference test (pLDDT) and predicted align error (PAE) (36, 66). These metrics respectively provide a per-residue accuracy metric of the generated model (pLDDT) and the expected positional error at residue *i* if the model is aligned on residue *j* (PAE) (36). The predicted RAD51-BRC4 complex was characterized by high pLDDT values, suggesting high confidence in the prediction output, except for flexible regions such as the DNA binding loops (L1 and L2) and the first 20 amino acids of the N-ter (**Figure S1A**). Moreover, we also observed low pLDDT values in the oligomerization linker region (amino-acids 86 to 89) where we substituted in the aminoacidic sequence F86 and A89 with E residues to match the aminoacidic composition of samples used for previous SAXS data collections on the RAD51-BRC4 complex (**Figure S1A**) (13). Interestingly, we also noted that PAE matrix reported a significant positional error of the relative position of RAD51 N-ter with respect to the RAD51 C-ter, possibly suggesting flexibility or complex conformational dynamics of this regions when it is not buried in another RAD51 monomer (**Figure S1B**, **S1C**). In the generated prediction the binding of BRC4 to RAD51’s C-ter matched the experimental structure of the complex, with a root mean square deviation (RMSD) computed on the Cα carbons of 4.0 Å (**Figure 2B**). Moreover, the N-ter was observed to preserve its internal folding (RMSD=0.7 Å), while its conformation significantly differed from the one found in RAD51’s *apo* structure, with an RMSD of 15.5 Å. In this way, BRC4 could bind without any clashes with the RAD51 N-ter (**Figure 2B, Figure S1C**). Interestingly, Alphafold2 predicted a network of polar contacts stabilizing the interaction of the BRC4 C-ter with the RAD51 N-ter, suggesting another important interface for the peptide binding (**Figure S1D**). To test the existence of this putative interaction between BRC4 and the protein N-ter, we decided to apply cross-linking mass spectrometry (XL-MS). Indeed, this technique would provide us useful structural information overcoming the difficulties of achieving a 3D crystal structure, as we observed the presence of different lysine residues (lys) at the predicted interface between the BRC4 peptide and the RAD51 N-ter (**Figure S1D**).

Considering the small size of the BRC4 peptide, we engineered a modified version harboring a biotin at the N-ter (bioBRC4) which we utilized to reconstitute *in vitro* the RAD51-BRC4 complex by mixing bioBRC4 with the monomeric His-RAD51[F86E, A89E] (hereafter referred to as “monomeric RAD51”). This would easily allow the identification of crosslinks by performing a Western Blot analysis tracking the bioBRC4 through Streptavidin coupled to Horse Radish Peroxidase (Streptavidin-HRP). We initially confirmed the binding of bioBRC4 to monomeric RAD51 by exploiting two orthogonal methods, Static Light Scattering (SLS) and Biolayer Interferometry (BLI) analyses (**Figure S2**).

Having assessed that the modification of the peptide did not negatively affect its binding to monomeric RAD51, we crosslinked the reconstituted bioBRC4-monomeric RAD51 complex, using both 1-ethyl-3-(3-dimethylaminopropyl)carbodiimide hydrochloride (EDAC) and bis(sulfosuccinimidyl)suberate (BS3) with spacer lengths of 0 and 11.4 Å, respectively. Initially, coomassie blue stained SDS-Page gels analyses suggested that BS3 crosslinked bioBRC4 to the monomeric RAD51 more effectively than EDAC. We then confirmed our observation by Western Blot analysis in which the bioBRC4 signal was clearly shifted to a molecular weight between 37 and 50 kDa matching a cross-linked RAD51-BRC4 complex (**Figure S3**). At this stage, we analyzed the cross-linked complex using LC-MS/MS on an Orbitrap Eclipse instrument. XL-MS analysis led to the identification of four high-quality crosslinks between BRC4 and RAD51 (**Figure 3A**, **S4 and Table 1**). Specifically, we observed that Lys1536 and Lys1541, in proximity of the BRC4 LFDE domain, were found to be crosslinked with Lys59, Lys65 and Lys71 located on the RAD51 N-ter in a region encompassing a cluster of alpha-helices, which support RAD51 interaction with the DNA (**Figure 3A**, **S4 and Table 1**) (67).

**Figure 3.**
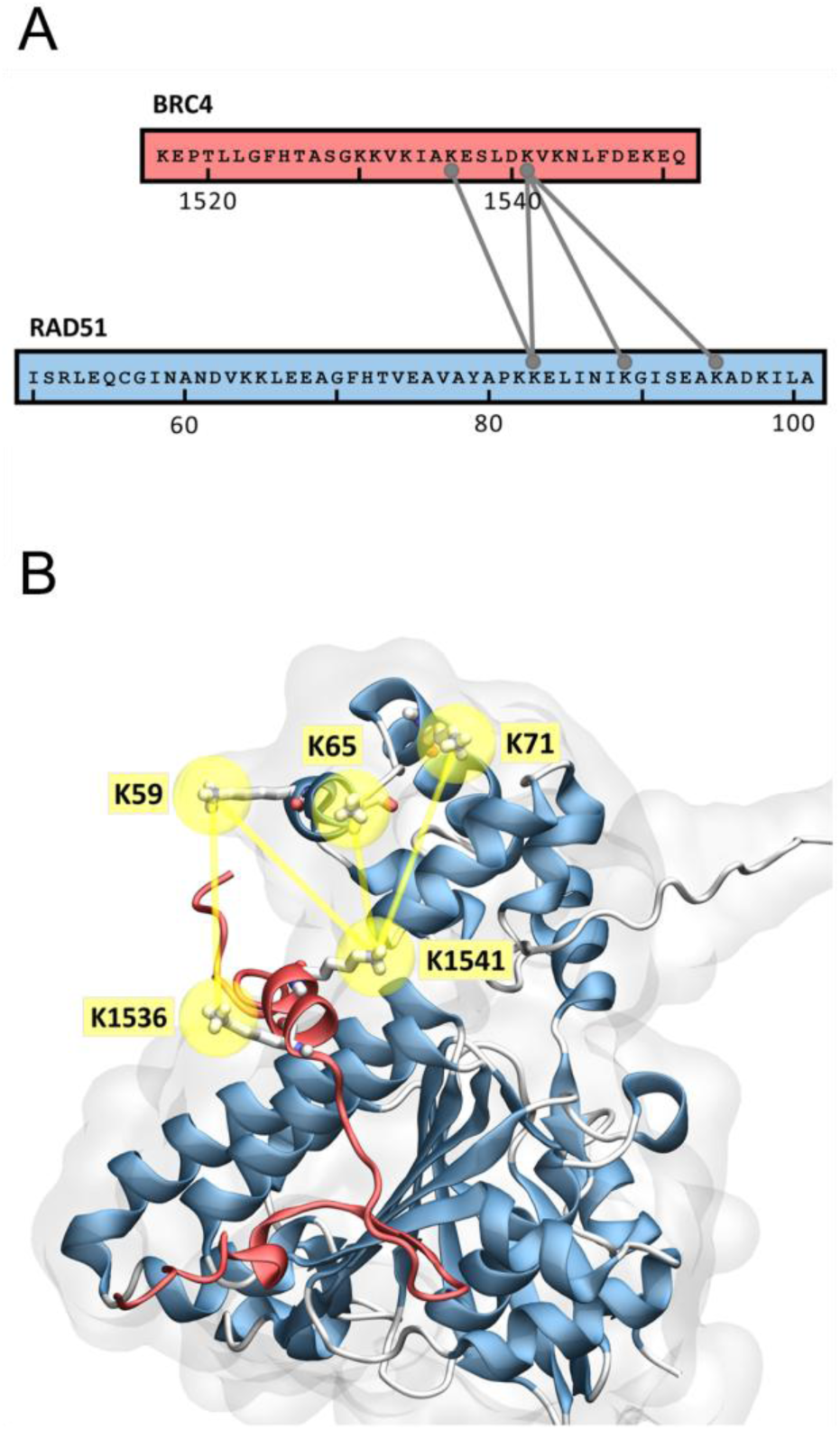
XL-MS data shows crosslinks between RAD51 and BRC4. A) Sequence overview of the BRC4 repeat of BRCA2 and RAD51 illustrating the detected inter-protein crosslinks filtered for MS quality score > 30 and Cα-Cα distances of 10 – 35 Å. Connecting lines point towards the cross-linked residues. B) Mapping of the identified crosslinks in the AlphaFold2 prediction of the RAD51-BRC4 complex.

**Table 1.**
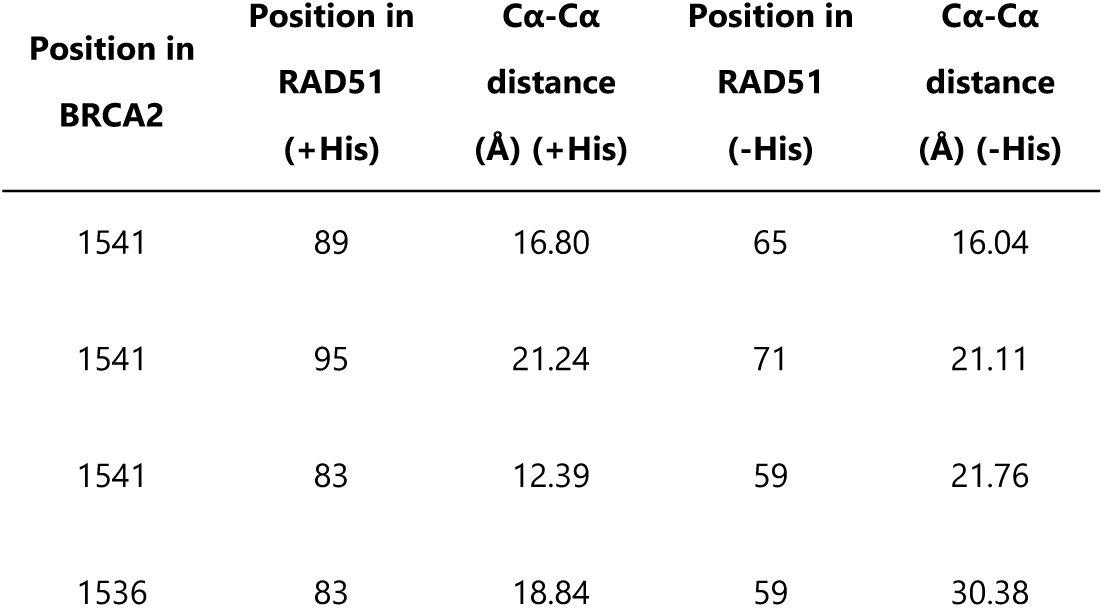
Table of detected inter-protein crosslinks and their measured Cα-Cα distances in the AlphaFold2 prediction of the His-RAD51 / BRC4 complex (+His) and the RAD51 / BRC4 complex (-His).

This result highlighted that the BRC4 C-terminus is crucial to allow the binding of the peptide to RAD51, since it directly interacts with the RAD51 N-ter, displacing it. Then, we aimed to verify the compliance to crosslinking data of His-RAD51/BRC4 and RAD51/BRC4 AlphaFold2 models (**Figure 2**, **Figure 3**, **Figure S1**, **Figure S5**, **Figure S6**). Notably, in both predictions the identified pairs of cross-linked lysine residues displayed Cα-Cα distance compatible with the experimental XL-MS results (**Table 1**) and within the limits of typical distance constraints, further corroborating our initial observation, made on the RAD51-BRC4 AlphaFold2 model, that a hydrogen bond network amidst the BRC4 C-terminus and the RAD51 N-term exists (53). Additional validation of the generated AlphaFold2 predictions was provided by distances of identified intra-RAD51 crosslinks. Indeed, the majority of identified crosslinks matched permissive distances in the predicted structure, with only two exhibiting long distances > 45 Å (**Figure S7**). While we cannot *a priori* exclude that the latter are experimental XL-MS false identifications, they may also derive from RAD51 N-ter flexibility in solution when BRC4 is bound (13). Indeed, the conformational rearrangements of this domain in solution could potentially bring the Lys residues in closer proximity to each other, thus allowing the crosslinking reaction by BS3.

### Including the solvent contribution is necessary for generating a SAXS-consistent single-structure model of the RAD51-BRC4 complex

Having validated the RAD51-BRC4 AlphaFold2 prediction by XL-MS, we then compared its computed spectrum with a recently determined SAXS spectrum for the RAD51-BRC4 complex in solution, already available in SASBDB (13). As **Figure 4A** shows (top panels), a significant discrepancy was observed, indicating that the initial guess, albeit reasonable from a merely structural standpoint and in terms of XL-MS data, was not representative of the RAD51-BRC4 complex in solution. Such difference can be further appreciated in the Kratky form of the spectra (**Figure 4A**, bottom panels), where the predicted SAXS profile suggested an overly compact configuration compared to the experimental one (68).

**Figure 4.**
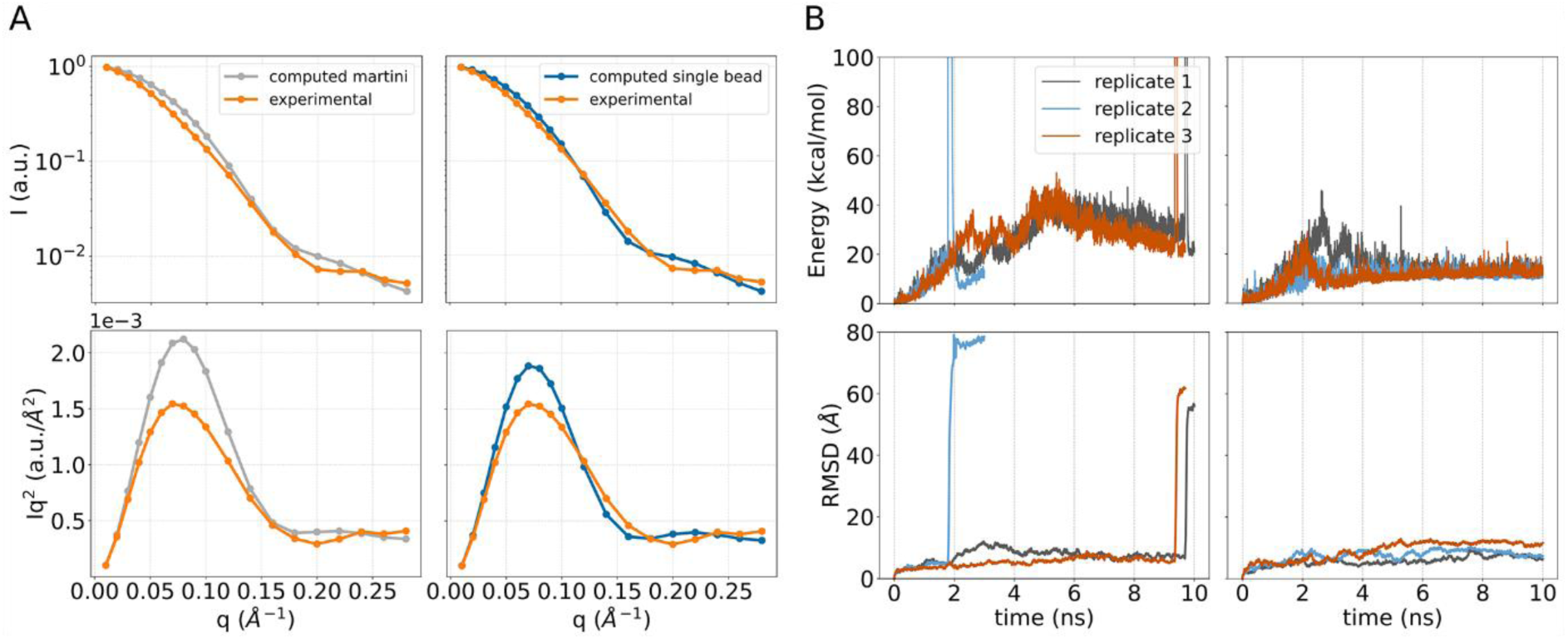
A) Logarithmic scale (top) and Kratky (bottom) plots of the SAXS spectrum computed on the AlphaFold model without (left) and with (right) solvent contribution, compared with experimentally measured SAXS spectrum of the complex. B) Timeseries of the sMD bias (top) and BRC4 heavy atoms RMSD after alignment on the C-ter (bottom), carried out without (left) and with (right) solvent contribution.

To guide the initial guess towards a configuration in better agreement with experimental data, we took advantage of steered MD (sMD) simulations. sMD is an enhanced sampling method where the exploration of complex biomolecular processes along a predefined CV is improved by using a time-dependent biasing potential. Typically, the biasing potential takes the form of a harmonic restraint that moves at a constant velocity during the simulation, driving the system towards a target value of the CV. Here, the system was driven towards the target state, defined by the experimental SAXS spectrum, through a hybrid all-atom/coarse-grain scheme (see Methods) over the course of 10 ns-long sMD simulations. Unexpectedly, in all three replicates, the simulation invariably resulted in the detachment of the BRC4 repeat from RAD51, as indicated by a rapid rise in the energy and quantified by a marked increase in the RMSD of the complex (**Figure 4B**, left top and bottom panels, respectively). Notably, this behavior is in contrast with the experimental SAXS data, which refer to the formed complex, as we have formerly demonstrated (13). We note that this set of calculations relied on a forward model that does not take into account the contribution of the protein’s solvation layer to the SAXS spectra (MT-hySAXS scheme). Remarkably, repeating the procedure by taking into account solvation effects (SB-hySAXS scheme) led to the preservation of the RAD51-BRC4 complex, as indicated by the absence of sudden rises in the energy as well as in RMSD (**Figure 4B**, right panels). Importantly, most of the conformational rearrangements required to match the experimental SAXS spectrum involved the N-ter (**Figures S8** and **S9**). We stress that this result was achieved by the simulations without providing any information about flexible regions in RAD51, nor by adding any restraint on the BRC4 repeat to disfavor detachment. Thus, in the effort of satisfying the experimental SAXS data, the system was able to naturally adapt based on its physical properties. We refer to the configurations attained by the system at the target state of the sMD simulation, obtained using the SB-hySAXS scheme, as single-structure models. Three replicates of sMD with the SB-hySAXS scheme resulted in comparable single structure models (**Figure S8** and **S9**). These models were instrumental for providing a SAXS-consistent initial state for the subsequent XL-MS-informed metadynamics simulations. Here, we used the single structure model from the first replicate of SB-hySAXS sMD for the subsequent stage.

### XL-MS-informed simulations enable determination of a SAXS-compliant conformational ensemble of the RAD51-BRC4 complex

As previously suggested, binding of the BRC4 peptide to RAD51 could trigger a conformational re-arrangement of the RAD51’s N-ter, which could behave as an intrinsically disordered domain (13). As a result, the marked conformational flexibility of this region makes the structural characterization of the FL-RAD51-BRC4 complex challenging (13). In such cases, an ensemble of conformations could better be suited to achieve a more realistic description of the system, rather than a single structure model (32, 69). In this respect, SAXS data become particularly useful since the measured spectrum results from the average over different conformations of the biomolecule in solution, becoming of remarkable importance to describe highly flexible molecules (e.g. IDPs, IDDs, RNAs). Given this scenario, we aimed to identify a conformational ensemble of structures that would be compatible with the experimental SAXS data (13). To this end, we first generated a heterogeneous ensemble of RAD51-BRC4 configurations, then the population weights were refined through a reweighting procedure in order to match experimental data. For the first stage, MD simulations did not include any information about the experimental SAXS spectrum, except for the use of the SAXS-consistent single-structure of the RAD51-BRC4 complex obtained from the first replicate of the sMD simulations as a starting configuration. Specifically, we carried out Metadynamics (metad) simulations, an enhanced sampling method that allows a comprehensive exploration of the configurational space along predefined CVs by applying a time-dependent Gaussian-shaped bias potential. Taking the inverse of the total bias potential at the end of the metad simulation enables the reconstruction of the corresponding free energy profile. In this case, to promote the exploration of RAD51-BRC4 configurations with different degrees of structural compactness, we used as a CV the radius of gyration of the complex (**Figure S10A** and **S10B**). Additionally, metad simulations were integrated with information from the XL-MS experimental data in the form of distance restraints between RAD51’s N-ter and the BRC4 repeat (**Figure S10C**). Including such information avoided sampling of irrelevant states that would be incompatible with the maximum distances as suggested by the XL-MS data. This favored a more effective exploration of the system’s configurational space, and, in turn, optimized the efficiency of the simulation. Therefore, the RAD51-BRC4 structures produced in this way were used as prior ensemble for the subsequent reweighting procedure according to the maximum entropy principle, using the experimental SAXS spectrum as ground truth. To avoid overfitting, we introduced a regularization term in the procedure, chosen to result in an optimal value of the reduced *χ*^2^ between reweighted and experimental SAXS spectra of about 1 (see Methods). As a result, we were able to identify an ensemble of structures with a calculated SAXS spectrum in agreement with the experimental one within statistical error (**Figure 5**, and **S11**). We note that this reweighting stage resulted in a Kish effective sample size of 7391, out of the 20001 structures used in the analysis. This indicates that a significant amount of structures from the prior ensemble, from the metad simulation, were retained in the maxent-reweighted one and effectively contributed to the final computed SAXS spectrum.

**Figure 5.**
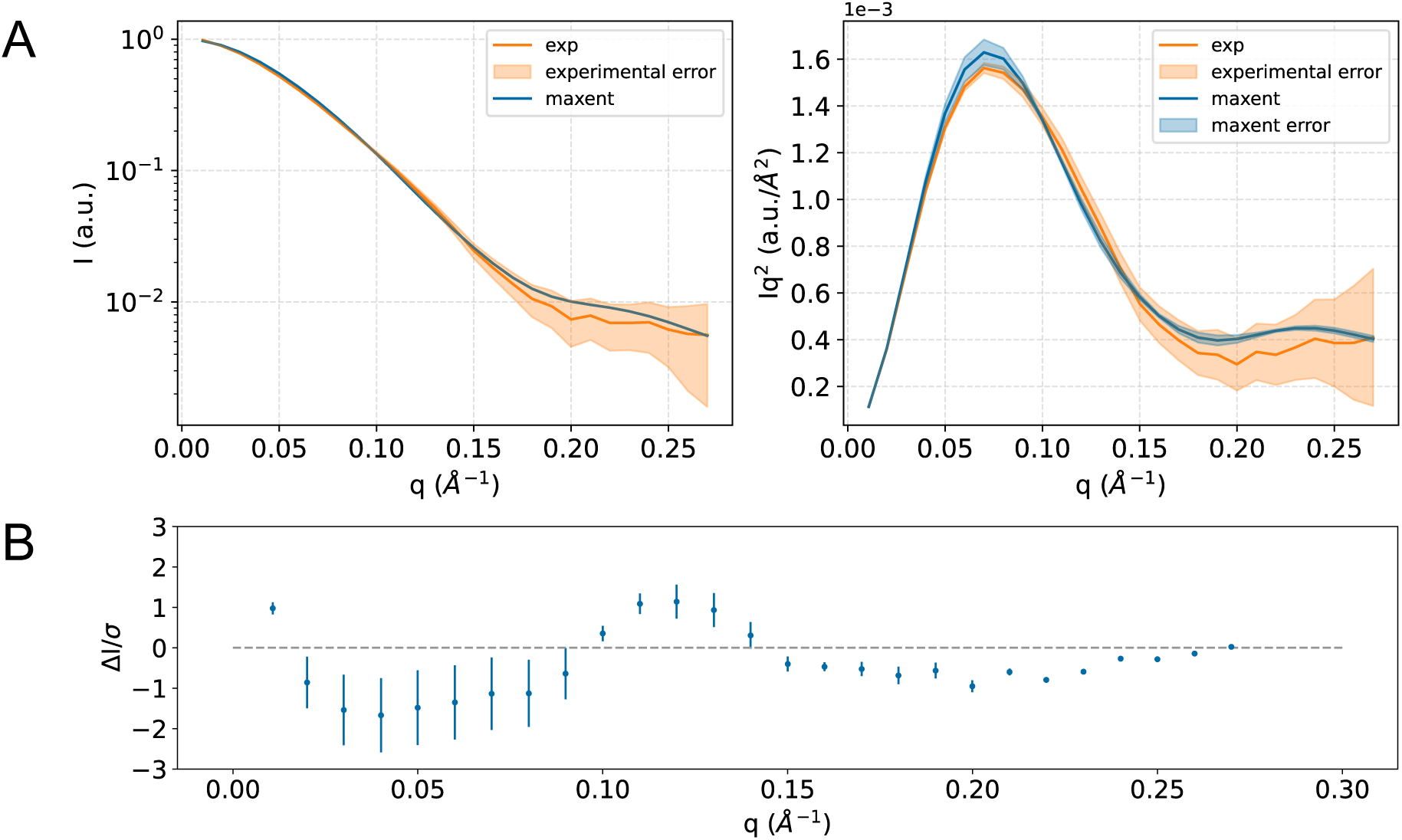
A) Comparison of the reweighted SAXS spectra through maxent with the experimental one, in logarithmic scale (left) and Kratky form (right), for the RAD51-BRC4 complex. B) Residuals plot of the computed spectrum with respect to the experimental one.

### Charged residues at the N-ter-BRC4 interface bridge compact and extended structures in the reweighted ensemble

Having a closer look at the reweighted ensemble allowed us to observe that both conformations with low values of the radius of gyration (compact), and higher ones (extended) were present. Interestingly, as shown in the reconstructed free energy profile (**Figure 6**), the prior ensemble (from metad) featured both compact structures within a deeper energy well centered at Rg ≈ 2.3 Å, and more extended ones, which however were considered less relevant. After maxent reweighing, higher weight was assigned to the more elongated structures in the shallow energy well centered at Rg ≈ 2.9 Å, resulting in basins of compact and extended structures having comparable depth (**Figure 6**). In particular, we observed that maxent reweighting allowed us to obtain a more balanced ensemble compared to the initial one obtained by metad. Indeed, after maxent the final population was constituted of a 65% of extended conformations while the remaining 35% was composed of compact structures. By contrast the metad population was almost totally composed of compact conformations (**∼**99%).

**Figure 6.**
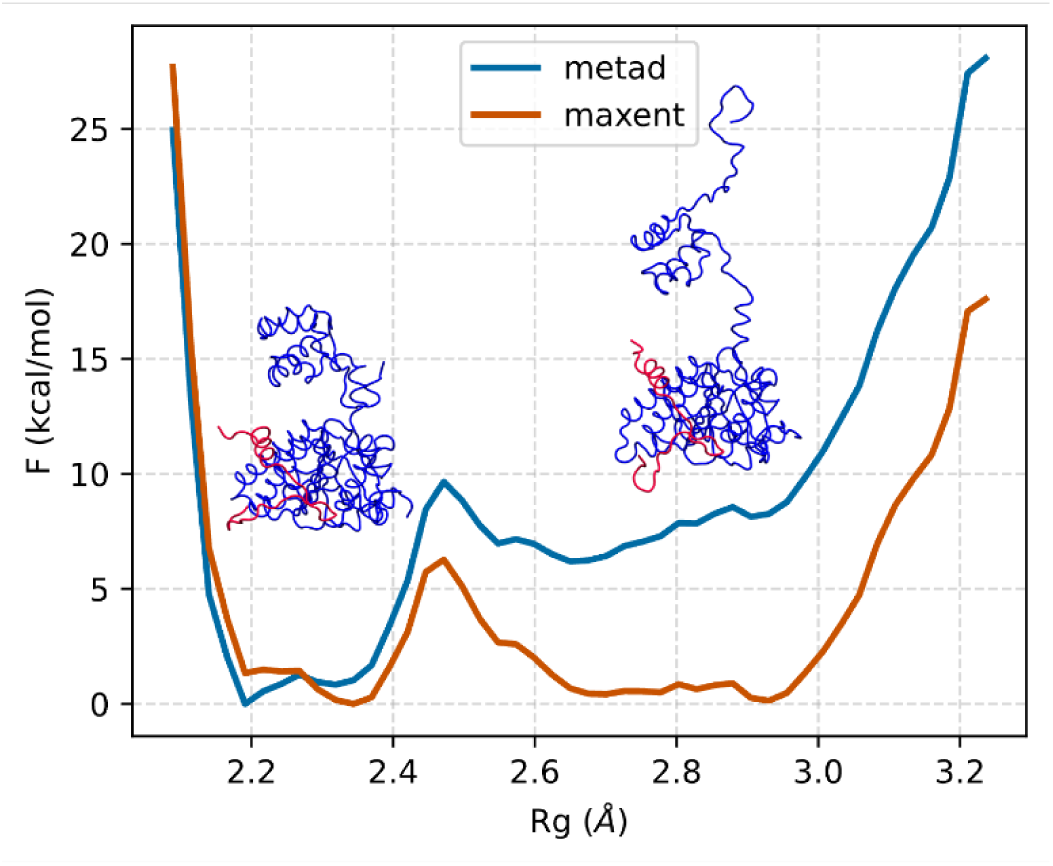
Free energy as a function of the radius of gyration for the metad ensemble (blue) and the reweighted ensemble via maxent (orange).

To better appreciate the observed maxent conformational heterogeneity of the RAD51-BRC4 complex in solution, we decided to represent it as a conformational space network as reported in **Figure 7**. Nodes in the network represent distinct clusters obtained from the metad simulations, with their size proportional to the corresponding maxent weights. The presence of edges indicates substantial similarity among different cluster representatives. Therefore, the network topology reflects the composition of the reconstructed conformational ensemble, allowing us to appreciate the distinctive features of the compact and extended subsets of structures (shown as orange and light blue nodes in **Figure 7**, respectively) and their mutual relationships.

**Figure 7.**
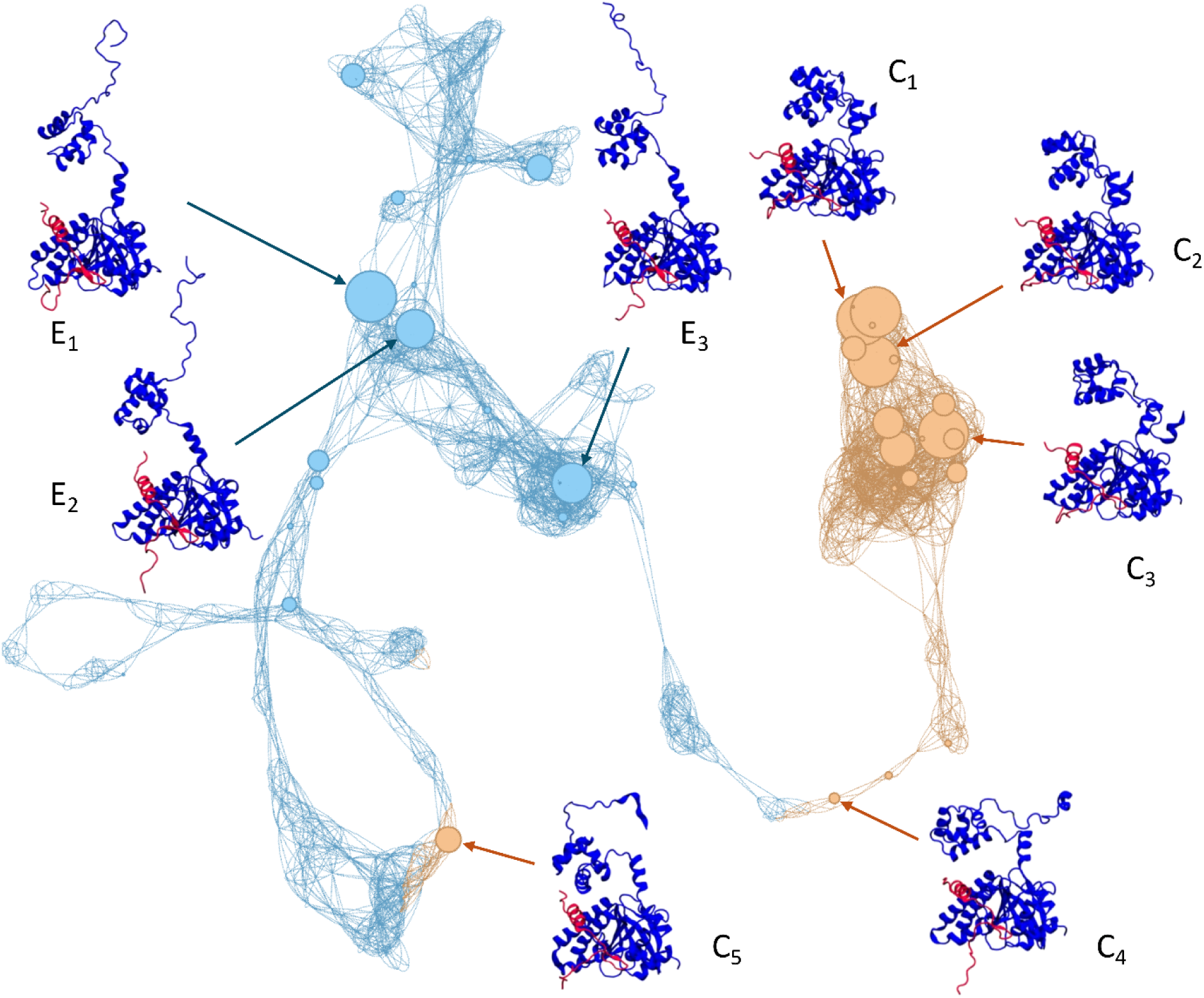
Conformational space network showing the reconstructed RAD51-BRC4 ensemble. Each node represents a cluster, whose size is proportional to the maxent reweighted population. Nodes are colored based on the degree of structural compactness, as indicated by the reweighted free energy profile in Figure 6 (orange for compact, light blue for extended). The representative structures of selected nodes, corresponding to either the most populated clusters or regions with peculiar network topology, are shown to highlight the conformational heterogeneity of the reconstructed ensemble.

Most compact structures are located in a dense, highly interconnected region of the plot, highlighting a strikingly conformational similarity among the representative members of different clusters. Nodes C_1_, C_2_, and C_3_ represent compact states with high statistical weight, showing an increasing and progressive separation between the RAD51’s N-ter and BRC4. Conversely, the regions of the network representing extended structures are more scattered, reflecting greater dissimilarity due to a higher conformational freedom associated to the full detachment of RAD51’s N-ter and increased disorder. Nodes E_1_, E_2_, and E_3_, are examples of clusters with high statistical weight belonging to the extended subset of conformations. Additionally, node C_4_ emerges as an intermediate structure connecting the compact and extended states, showing a substantially compact conformation with a partly disordered RAD51’s N-ter. While marginal in statistical weight, isolated compact states also appear in different regions of the network, with node C_5_ being the most populated. As revealed by the conformational space network, the difference in structural compactness is mainly governed by configurational rearrangements of the N-ter domain with respect to the complex between BRC4 and RAD51’s C-ter (**Figure 7**).

To gain further insights into the mechanistic details regulating the RAD51-BRC4 complex formation, we analyzed the interaction at the interface between BRC4 and RAD51’s N-ter (**Figure 8**, **Figure S12**). In particular, a contact analysis of the structures in the reweighted ensemble revealed that the most frequent contacts feature charged aminoacidic residues, namely lysine and glutamate. Specifically, Lys59 from the N-ter and Glu38 and Gln39 from the BRC4 repeat were mainly involved in the interaction between BRC4 and the N-ter domain.

**Figure 8.**
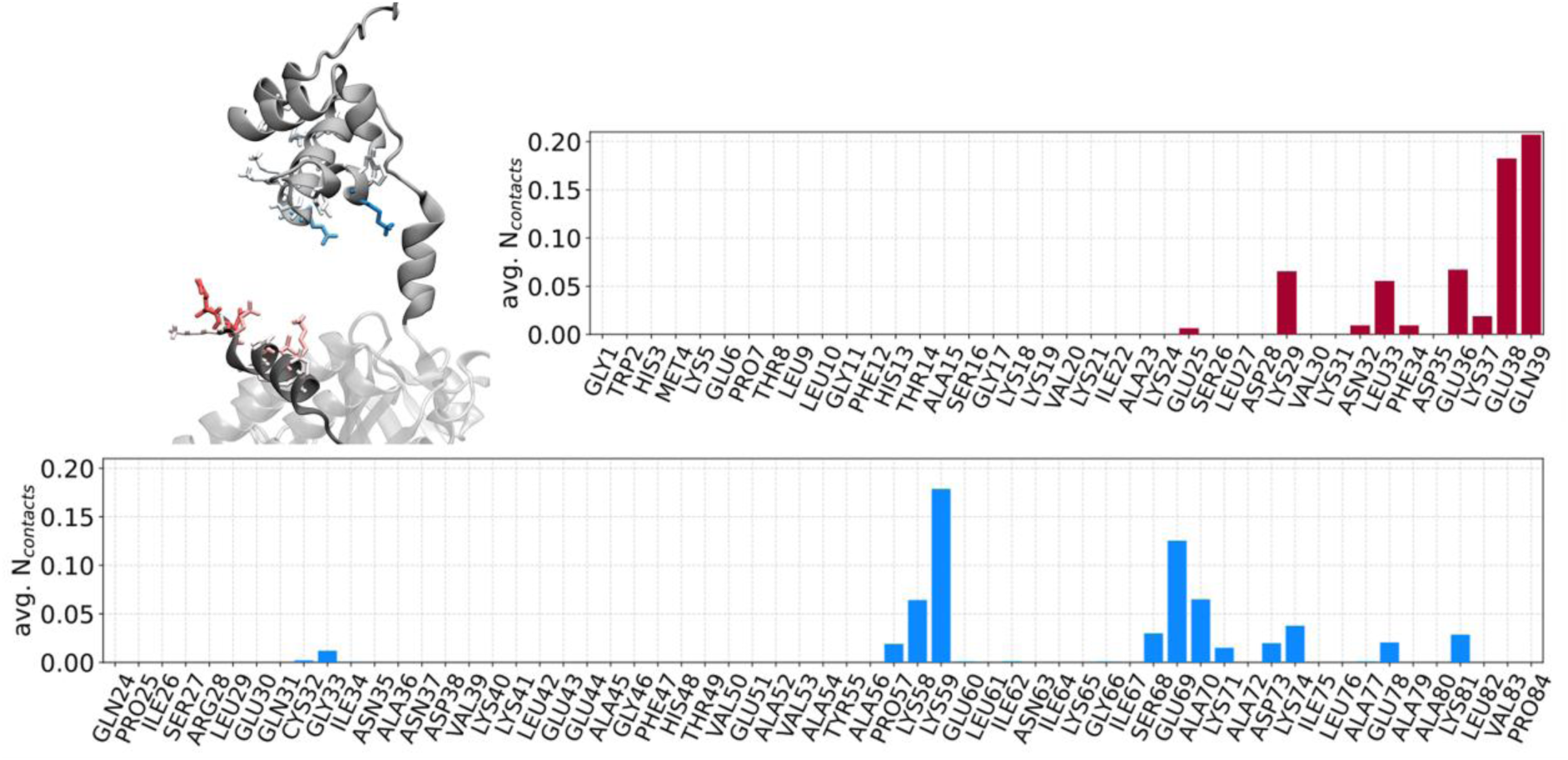
Residue-wise average number of contacts for the structures in the maxent-reweighted ensemble. The analysis focused on contacts for residues the BRC4 repeat (in blue) and RAD51’s N-ter (in orange). Residues with non-zero contacts are displayed in licorice, colored consistently with the plots and with color ranging from white to full solid, indicating low to maximum number of observed contacts, respectively; the C-ter, N-ter and BRC4 are displayed in VMD’s NewCartoon representation, colored white, grey and dark grey, respectively.

## DISCUSSION

Structural information is essential to understanding the physiological functioning of biomolecules and how pathological dysregulations arise. Moreover, this information can provide valuable insights to elaborate rational strategies for novel therapeutic interventions. The integration of computational methods with experimental data is increasingly recognized as a powerful approach to characterize systems with increasing structural complexity, where computational or experimental methods alone would not prove sufficient. In this work, we aimed at reconstructing the conformational ensemble of the full RAD51 protein in complex with the BRC4 peptide, for which an experimental structure is still missing. To address the challenges posed by the high flexibility of RAD51’s N-ter domain, we have complemented advanced MD simulations with newly acquired XL-MS and recently published SAXS data.

XL-MS studies enabled us to test the existence of the interaction between the BRC4 C-terminus and the RAD51 N-terminal domain, thus providing initial validation of the predicted Alphafold2 model and valuable constraints for successive computational studies. Moreover, this result also sheds light on an important mechanistic insight in the BRC4 interaction with RAD51 fibrils. Indeed, once BRC4 binds to RAD51, its C-terminus destabilizes the interaction of the RAD51 N-ter with another monomer thus promoting its detachment from the fibrils (14). Then, we integrated SAXS data into sMD simulations through a hybrid all-atom/coarse-grain scheme to refine the model obtained through AlphaFold2. Interestingly, while this structure displayed distances between lysine pairs compliant with the XL-MS experimental data, it was incompatible with the experimental SAXS spectrum, regardless of the forward model used. Nevertheless, including the solvation layer contribution resulted in a predicted spectrum closer to the experimental one, underscoring its significant role in SAXS calculations for this system. This finding was further confirmed by the sMD simulations, where we observed that accounting for the solvent was critical to successfully drive the structure towards agreement with the experimental SAXS spectrum without causing the detachment of the BRC4 peptide from RAD51.

The resulting single-structure model, now consistent with the experimental SAXS spectrum, proved instrumental in generating reasonable prior ensemble via subsequent metad simulations, enriched with configurations compatible with experimental data. Additionally, incorporating XL-MS restraints in the metad simulations discouraged the exploration of less relevant states, thus increasing the efficiency of the simulations and optimizing the exploitation of the computational resources. Notably, this initial ensemble of structures, which spanned a wide range of structural compactness, provided a suitable prior for successful reweighting via the maximum entropy principle. This was reflected by the obtained Kish size, indicating that a significant fraction of structures from the prior contributed to the reweighted ensemble. Specifically, a larger proportion of extended structures needed to be included and assigned higher weights to achieve a reconstructed ensemble in agreement with the SAXS experimental data, raising their population from less than 1% to about 65%. Besides demonstrating that our sampling strategy was able to produce a prior ensemble of high quality, this remarked the feasibility of generating a reasonable prior resorting to MD simulation timescales that are currently within reach of most research groups.

To provide further insights into the RAD51-BRC4 interaction, we examined the reweighted ensemble from a mechanistic standpoint. Given the peculiar structural flexibility of the N-ter domain in defining the conformational dynamics of the RAD51-BRC4 complex after BRC4 binding, we prompted our focus on possible interactions between residues from the N-ter domain and the BRC4 repeat. We performed a contact analysis leveraging a rather permissive cutoff, to possibly identify residues mediating the transition between compact and extended configurations of the complex. This analysis pinpointed charged residues at the N-ter-BRC4 interface, particularly Lys59 from the N-ter and Glu38 and Gln39 from the BRC4 repeat. We note that such information was retrieved in a dynamical setting, as the structures have been identified from MD simulations. This suggests how the long-range nature of the interaction between charged residues may bridge the transition from compact to more extended states. Besides providing additional details in the comprehension of RAD51-BRC recognition, this information may be leveraged for the rational design of binders at the BRC4 site modulating RAD51’s activity.

Achieving agreement between experiments and simulations has been a long-standing goal in biophysics and structural biology. In this work, we demonstrate the importance of including SAXS and XL-MS data in MD-based simulations to obtain a reliable structural ensemble of the RAD51-BRC4 complex. While the initial AlphaFold2 model provided a reasonable guess of the complex, as confirmed by its agreement with XL-MS data, the inherent limitations of a static modeling approach prevented it from reproducing the SAXS spectrum of RAD51-BRC4 in solution. Nonetheless, the AlphaFold2 model was instrumental, given the presence of extensive unstructured regions, as it provided a reasonable starting point for the subsequent MD-based simulation.

To the best of our knowledge, this is the first report comprehensively characterizing the full conformational ensemble of the RAD51-BRC4 interaction in solution with atomistic resolution, gaining insights into critical residues underpinning their complex conformational dynamics. Our study further corroborates our initial hypothesis that the ability of BRC4 to depolymerize RAD51 fibrils is mainly related to the ability of the C-terminus of the BRC4 peptide to directly interact with the RAD51 N-terminal domain, which can then explore multiple conformations in solution. It is likely that this multifaceted behavior of the RAD51 N-ter importantly impairs its capacity to oligomerize and allows its recruitment by BRCA2 and the RAD51 translocation inside the nucleus (14). Dissecting the mechanistic details of the interaction with the BRC4 repeat is fundamental to understanding the molecular features governing the recognition between RAD51 and the BRCA2 protein, underlying severe pathological conditions associated with DNA damage repair. In this perspective, our study provides robust insights that can be leveraged for the development of rational strategies for therapeutic intervention targeting the BRCA2-RAD51 interaction.

## DATA AVAILABILITY

SAXS data supporting this study are openly available in SASBDB at https://www.sasbdb.org/, reference number SASDQT9 – Monomeric DNA repair protein RAD51 homolog 1 double mutant [F86E, A89E] in complex with fourth BRC repeat (BRC4) (70). The mass spectrometry proteomics data have been deposited to the ProteomeXchange Consortium via the PRIDE partner repository with the dataset identifier PXD060867 (71). We freely provide the input files (initial coordinates, topologies, GROMACS mdp parameter file, and PLUMED inputs) to perform the MD simulations in this work, as well as the output simulation trajectories (in xtc format) that we generated. We also supply a Jupyter notebook to reproduce all of our analyses, results, and the plots reported in this work. The material is freely available in Zenodo with accession code 10.5281/zenodo.14988991. The Jupyter notebook can be straightforwardly consulted and downloaded at github.com/CompMedChemLab/project_saxs-xlms-md_rad. All MD simulations were performed with GROMACS 2023.1, patched with plumed 2.9. The VMD software version 1.9.4 was used for visualization, and analyses were conducted with MDtraj version 1.9.9, MDAnalysis Version 2.6.1, scikit-learn version 1.3.1.

## SUPPLEMENTARY DATA

Supplementary Data are available at NAR online.

## ACKNOWLEDGEMENTS

The authors gratefully thank the European Institute of Oncology Biochemistry and Structural Biology Unit for useful discussions. We gratefully acknowledge the Data Science and Computation Facility and its Support Team at Fondazione Istituto Italiano di Tecnologia for computing time and support on the Franklin HPC system. We would also like to thank Imke Wüllenweber for excellent technical assistance for proteomics sample preparation. The authors thank the B21 at Diamond Light Source (DLS) for providing the synchrotron radiation facility for SAXS measurements and for fruitful discussions.

