## Supplemental Figure S1-S12 for "Integrating computational and experimental biophysics reveals novel insights into the RAD51-BRC4 interaction"

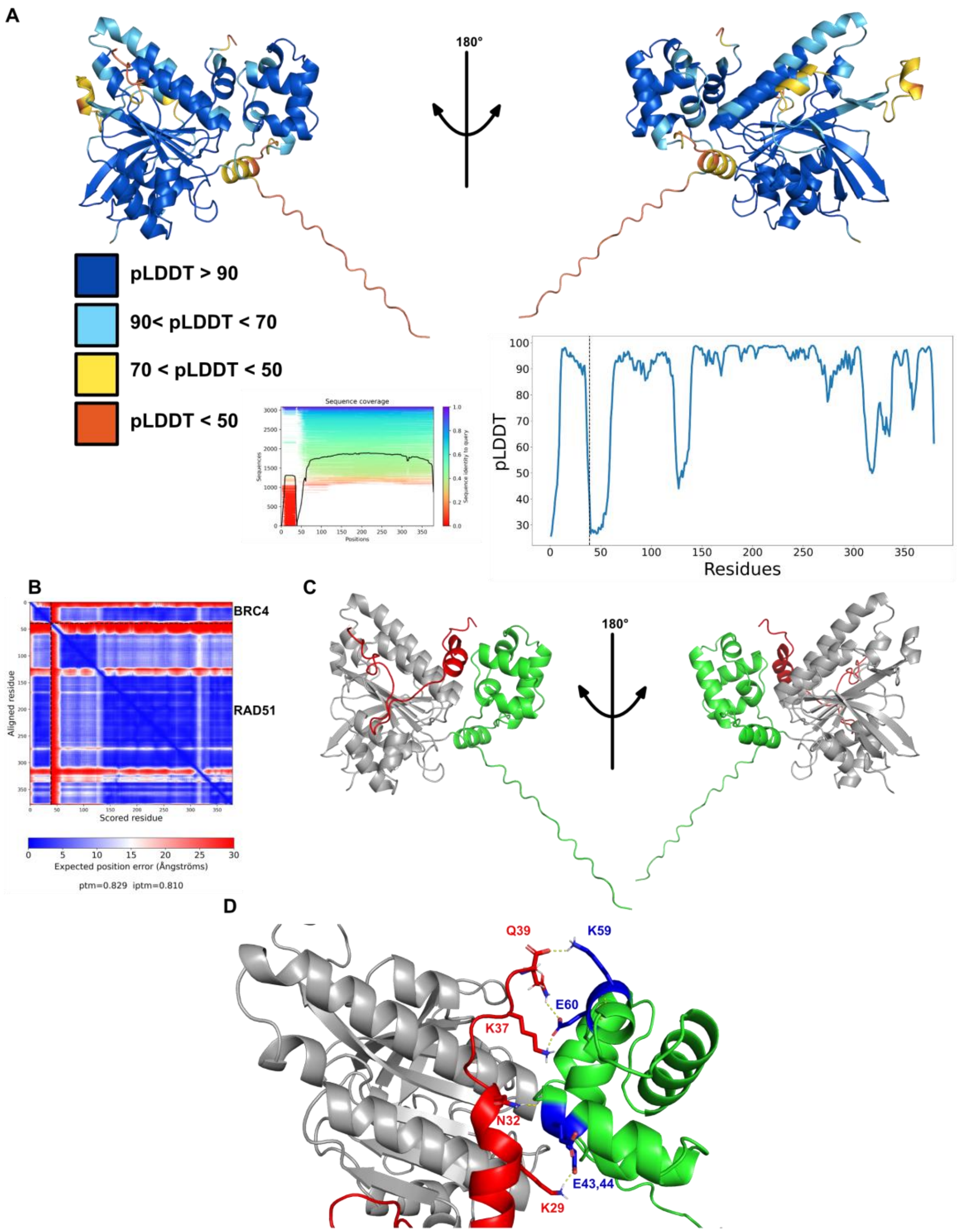


**Figure S1 Alphafold 2.3 model of full length RAD51 monomer in complex with BRC4. A.** Residues have been colored for local model confidence estimated with Predicted Local Distance Difference Test (pLDDT) representation (per-residue accuracy metric of the generated model) **B**. Predicted Align Error (PAE) plot, showing the expected positional error at residue x if the model is aligned on residue y. Right: Alphafold 2.3 Multiple Sequence Alignment. **C.** Residues have been colored by domain (green: N-terminal domain, grey: C-terminal domain) **D.** Analysis of polar interactions (hydrogen bonds) taking place between BRC4 and the RAD51 N-terminal domain predicted by Alphafold 2.3.

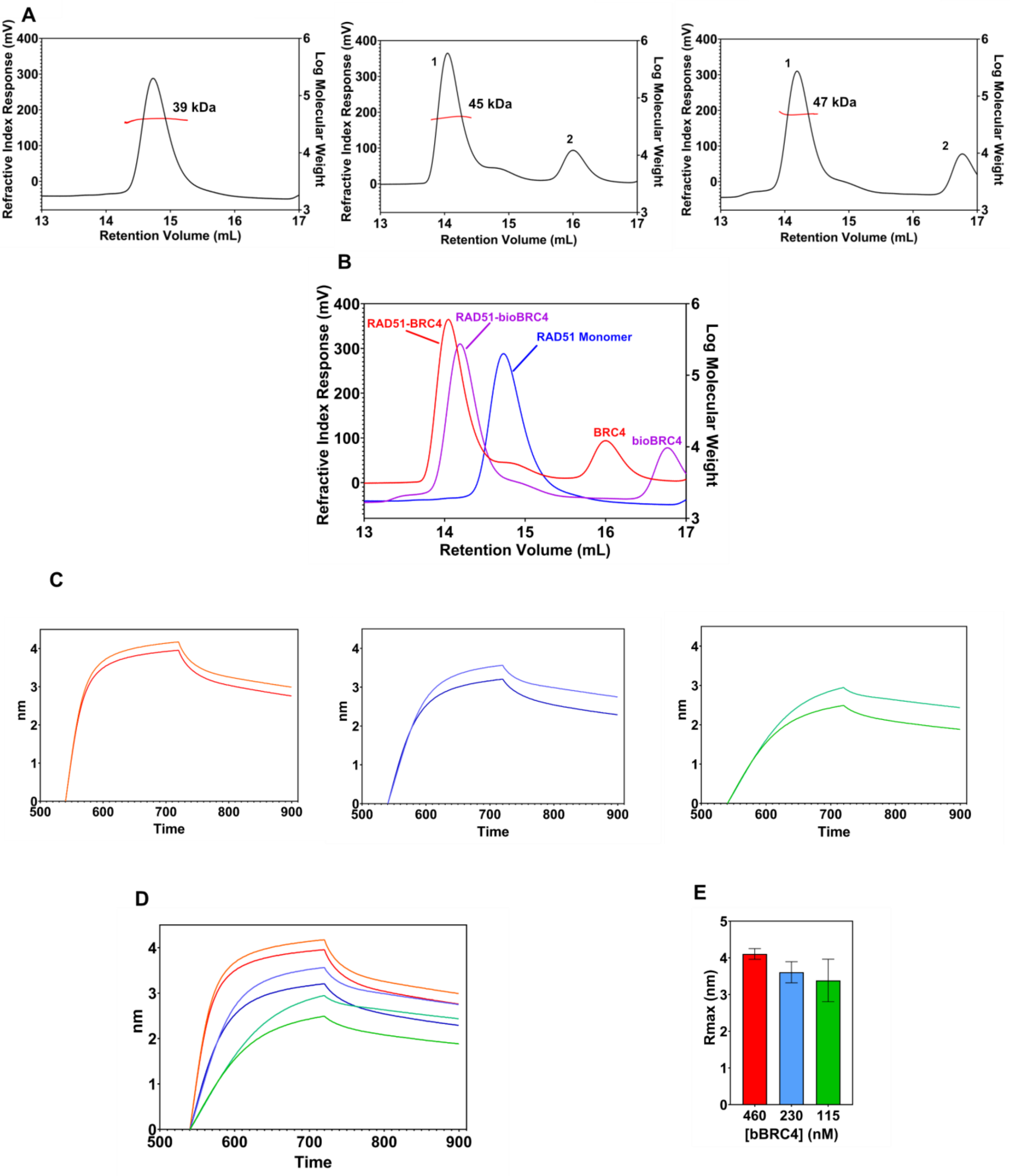


**Figure S2 Binding tests of the biotinylated BRC4 peptide (bio-BRC4) to the monomeric His-RAD51 [F86E, A89E]** **A.** SLS analysis in buffer containing 100 mM $Na_2SO_4$. Left: RAD51 [F86E, A89E] Middle: RAD51 [F86E, A89E] in presence of BRC4 peptide, peak 1 represents the RAD51-BRC4 complex, peak 2 represents the BRC4 peptide alone. Right: RAD51 [F86E, A89E] in presence of bioBRC4 peptide, peak 1 represents the RAD51-BRC4 complex, peak 2 represents the bioBRC4 peptide alone BRC4 peptide (green). **B.** Overlay view of the samples in the absence (blue) or presence of BRC4 (red) and bioBRC4 peptide (purple). **C.** Two replicates of biolayer interferometry (BLI)sensorgrams showing the binding of monomeric His-RAD51[F86E, A89E] to bioBRC4 obtained at three different protein concentrations. Left: 460 nM, Middle: 230 nM, Left: 115 nM **D.** Overlay of all BLI sensorgrams **E.** Mean ± standard deviation of maximum response ($R_{max}$) obtained for BLI experiments carried out at the same protein concentration

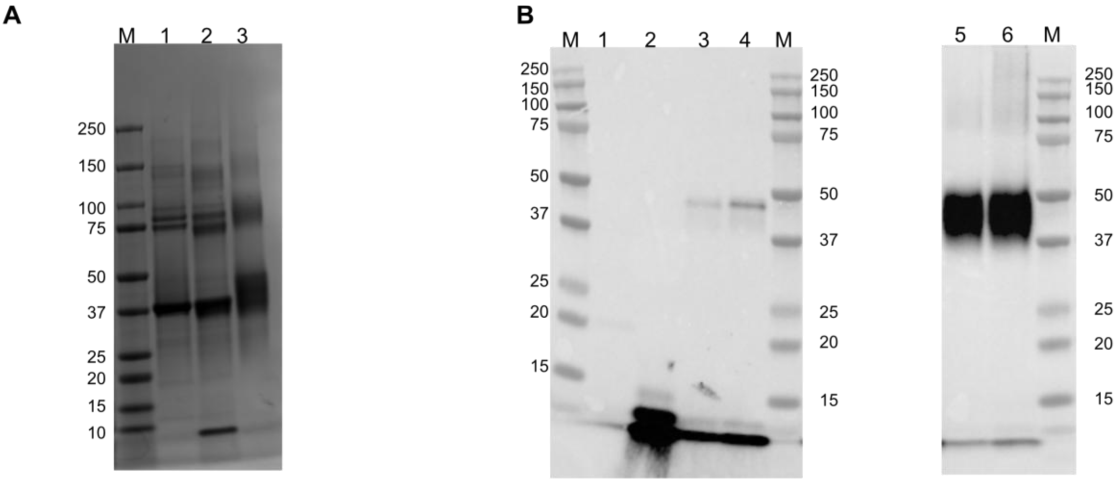


**Figure S3 Monomeric RAD51 [F86E, A89E] bioBRC4 cross-linking trials. A.** SDS Page gel Coomassie Blue Staining, M = Marker, 1 = RAD51 [F86E, A89E] 2 = RAD51:bioBRC4 complex 1:2 crosslinked with EDAC 0.2 % (w/v) 3 = RAD51:bioBRC4 complex 1:1 crosslinked with BS3 (1 mM) **B.** Western Blot detecting bio-BRC4 with Streptavidin-HRP M = Marker, 1 = RAD51 [F86E, A89E], 2 = bioBRC4, 3 = RAD51:bioBRC4 complex 1:1 4 = RAD51:bioBRC4 complex 1:2 5 = RAD51:bioBRC4 complex 1:1 BS3 1 mM 6 = RAD51:bioBRC4 complex 1:2 BS3 1 mM

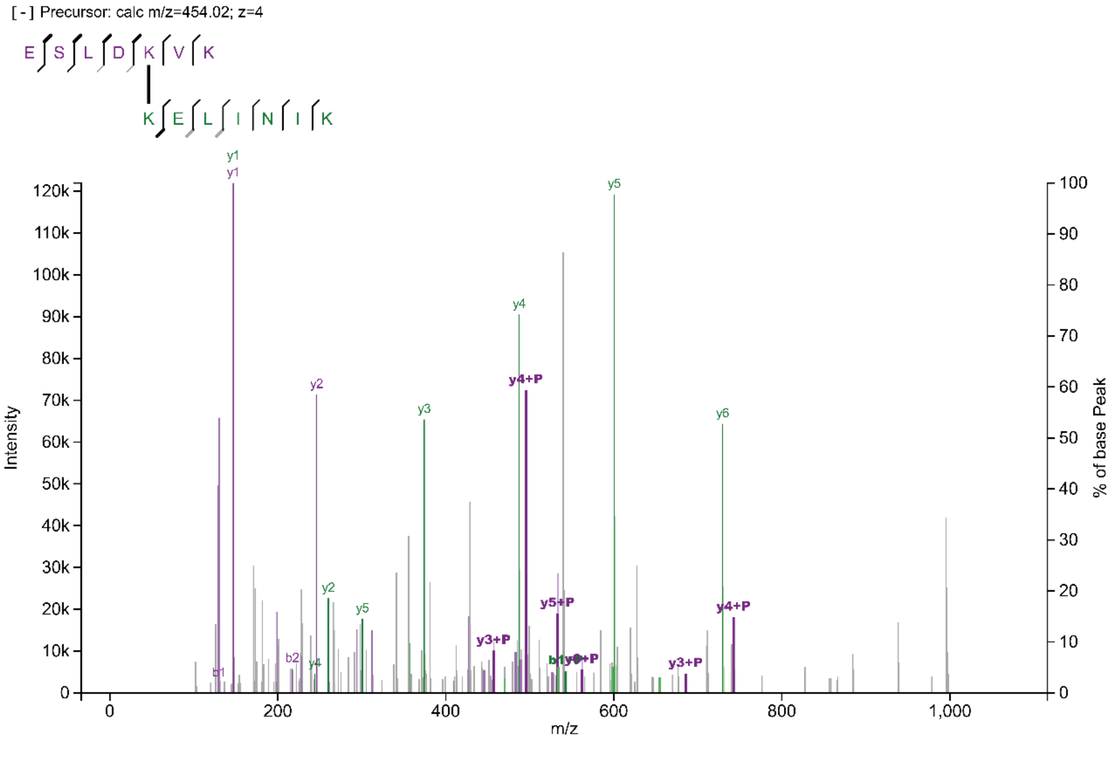


**Figure S4 Example mass spectra of crosslinked peptides, evidencing the quality of the identifications.**

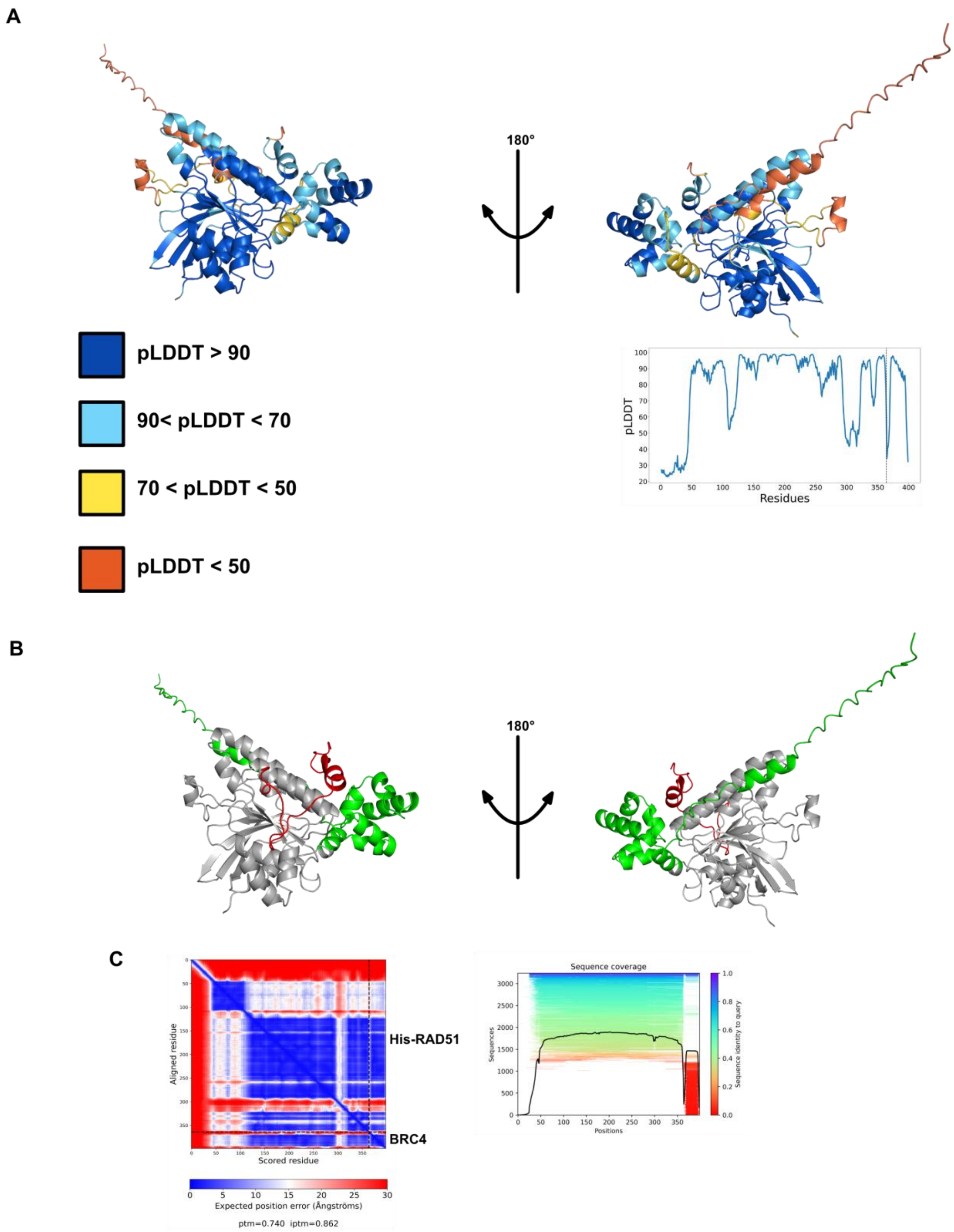


**Figure S5 Alphafold 2.3 model of His-tagged full length RAD51 monomer in complex with BRC4**. **A.** Residues have been colored for local model confidence estimated with Predicted Local Distance Difference Test (pLDDT) representation (per-residue accuracy metric of the generated model) **B.** Residues have been colored by domain (green: N-terminal domain, grey: C-terminal domain) **C.** Left: Predicted Align Error (PAE) plot, showing the expected positional error at residue x if the model is aligned on residue y. Right: Alphafold 2.3 Multiple Sequence Alignment.

**A**

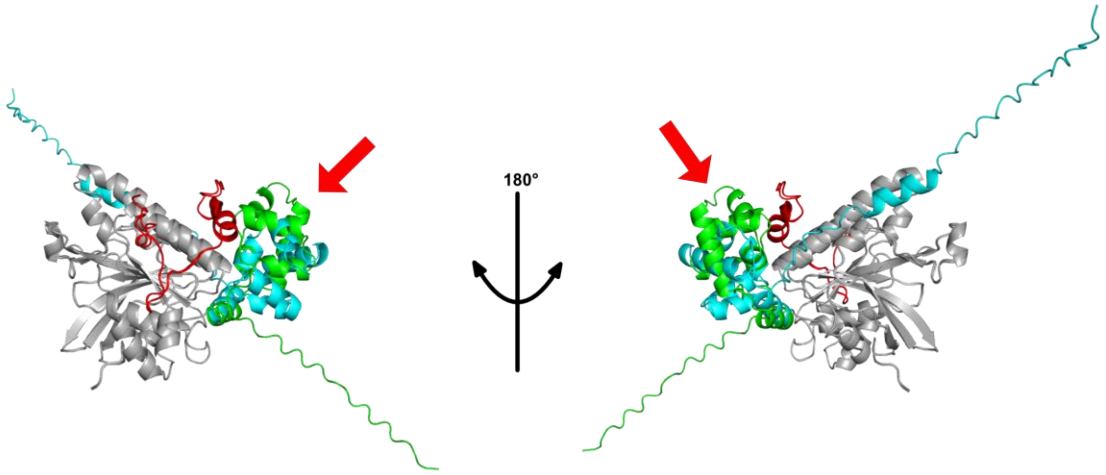


**R.M.S.D. = 0.303**

**B**

**RAD51-BRC4**

**>A**

GWHMKEPTLLGFHTASGKKVKIAKESLDKVKNLFDEKEQ

**>B**

GMAMQMQLEANADTSVEEESFGPQPISRLEQCGINANDVKKLEEAGFHTVEAVAYAPKKELINIKGISEAKADKIL
AEAAKLVPMGETTETEFHQRRSEIIQITTGSKELDKLLQGGIETGSITEMFGEFRTGKTQICHTLAVTCQLPIDRGG
GEGKAMYIDTEGTFRPERLLAVAERYGLSGSDVLDNVAYARAFNTDHQTQLLYQASAMMVESRYALLIVDSATAL
YRTDYSGRGELSARQMHLARFLRMLLRLADEFGVAVVITNQVVAQVDGAAMFAADPKKPIGGNIIAHASTTRLYL
RKGRGETRICKIYDSPCLPEAEAMFAINADGVGDAKD

**HisRAD51-BRC4**

**>A**

KEPTLLGFHTASGKKVKIAKESLDKVKNLFDEKEQ

**>B**

MGSSHHHHHHSSGLVPRGSHMLEDPMAMQMQLEANADTSVEEESFGPQPISRLEQCGINANDVKKLEEAGFH
TVEAVAYAPKKELINIKGISEAKADKILAEAAKLVPMGETTETEFHQRRSEIIQITTGSKELDKLLQGGIETGSITEMF
GEFRTGKTQICHTLAVTCQLPIDRGGGEGKAMYIDTEGTFRPERLLAVAERYGLSGSDVLDNVAYARAFNTDHQT
QLLYQASAMMVESRYALLIVDSATALYRTDYSGRGELSARQMHLARFLRMLLRLADEFGVAVVITNQVVAQVDGA
AMFAADPKKPIGGNIIAHASTTRLYLRKGRGETRICKIYDSPCLPEAEAMFAINADGVGDAKD

**Figure S6 Overlay of the Alphafold 2.3 generated models.** Overlay of the RAD51-BRC4 and of the His-RAD51-BRC4 complexes. Red arrows highlight the different arrangements of the RAD51 N-terminal domain in the two predicted models. In multiple colours different domains of RAD51 (green (RAD51-BRC4) and cyan (His-RAD51-BRC4) represent the RAD51 N-terminal domain while the C-terminal is displayed in grey) in the generated AF2 model and the BRC4 peptide (in red). The root mean square deviation (R.M.S.D.) of the two models is displayed in the figure. **B.** Amino acidic sequence of the generated AF2 models.

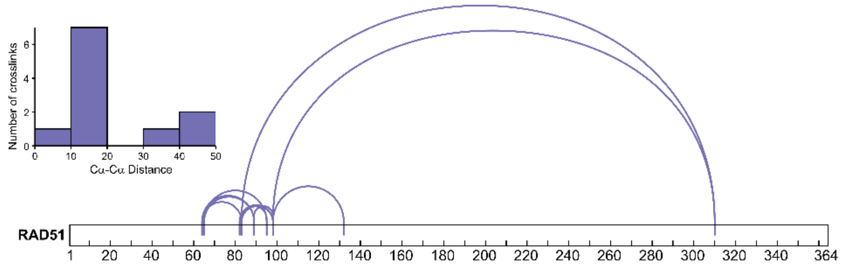


| Crosslink | Cα-Cα distance -His Tag | Cα-Cα distance +His Tag |
|---|---|---|
| K64 - K83 | 18.24 | 18.33 |
| K64 - K89 | 11.31 | 11.30 |
| K65 - K89 | 7.59 | 7.61 |
| K65 - K95 | 11.57 | 11.69 |
| K82 - K98 | 10.60 | 10.44 |
| K83 - K310 | 39.16 | 46.75 |
| K83 - K95 | 11.89 | 11.80 |
| K83 - K98 | 13.63 | 13.45 |
| K89 - K98 | 15.91 | 15.96 |
| K98 - K132 | 47.96 | 41.67 |
| K98 - K310 | 43.96 | 48.69 |

**Figure S7 RAD51 intramolecular cross-links identified in XL-MS analysis**

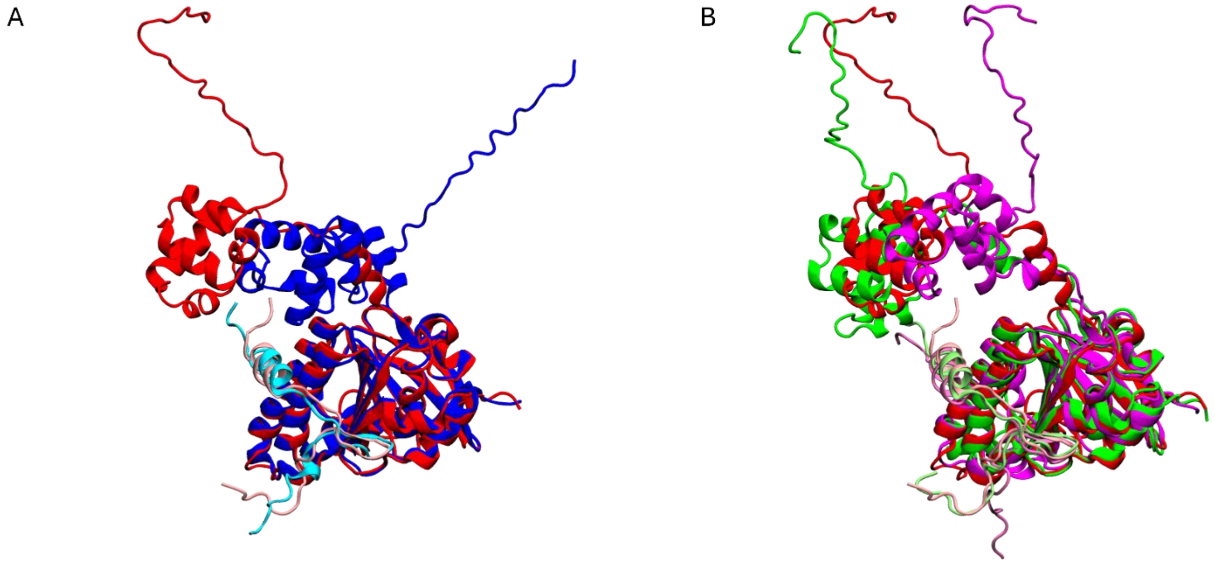


**Figure S8** A) One model from the single structure model from replicate 1 of the steered MD simulations via the SB-hySAXS scheme (red) superposed to the AlphaFold model (blue), and B) single structure models from all the three replicates of steered MD (red, green, and purple for replicates 1, 2, and 3, respectively). All structures are aligned on the heavy atoms of RAD51's C-ter domain.

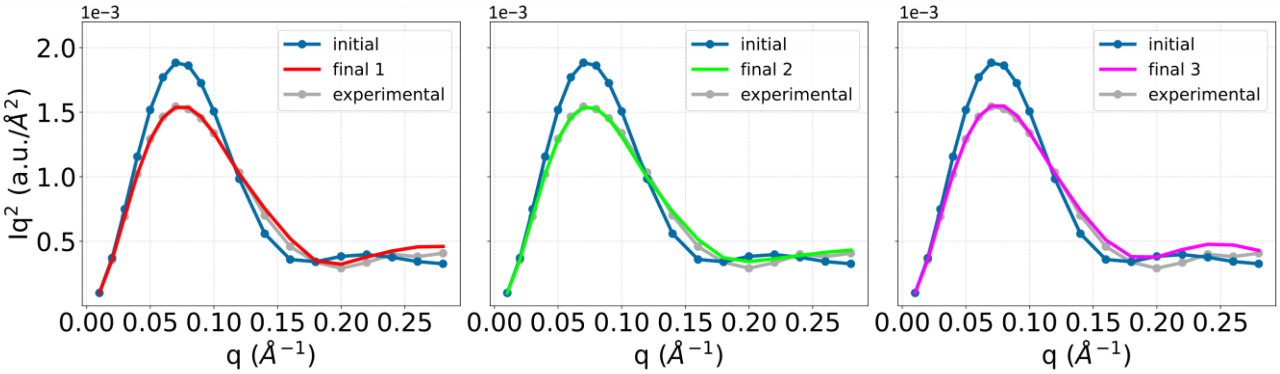


**Figure S9** SAXS spectra in the Kratky form for the three replicates of steered MD using the SB-hySAXS scheme. The three panels report the spectrum computed at the end of the simulation (labelled final, and color coded according to figure S3B), compared with the starting structure used for the simulations, i.e. the AlphaFold model (in blue), and the experimental SAXS spectrum (in grey).

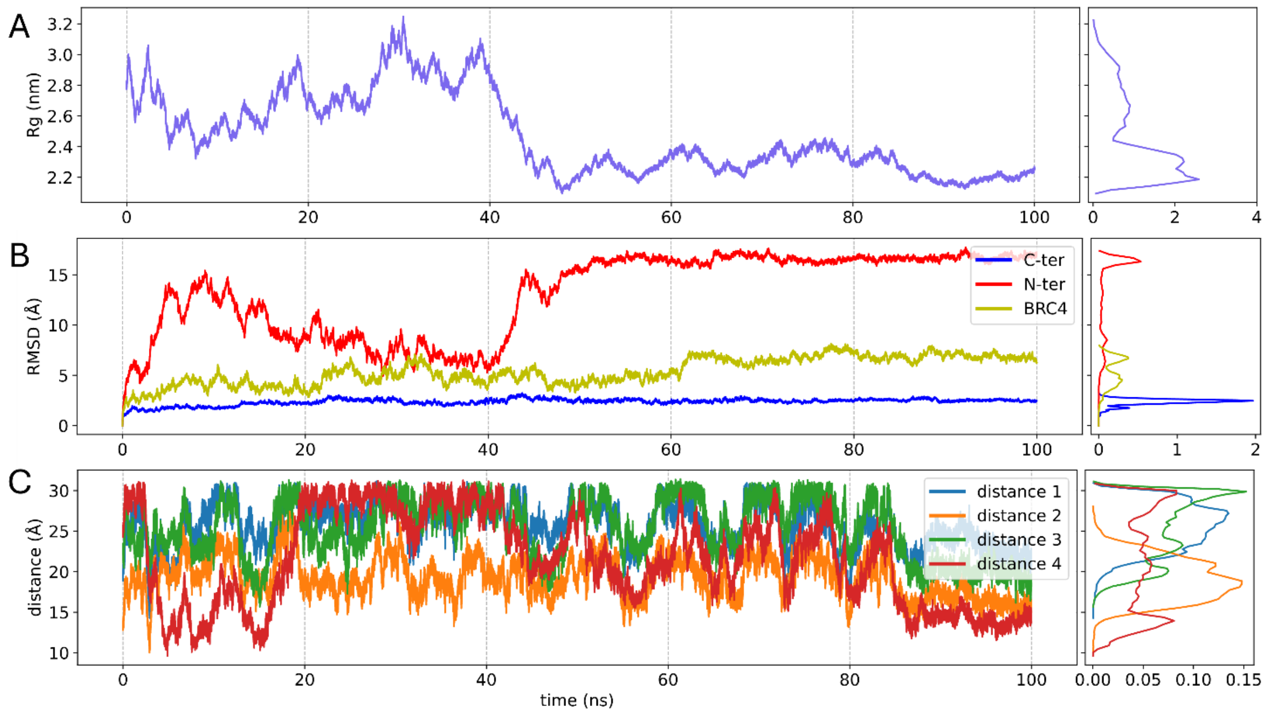


**Figure S10** Timeseries and corresponding distributions along the metad simulation for A) the radius of gyration of the system, B) RMSDs of RAD51's N-ter and C-ter, and the BRC4 repeat, and C) distances between lysine pairs from the XL-MS experiments. RMSDs were computed on heavy atoms, after aligning separately on each of the three characters (N-ter, C-ter, and BRC4).

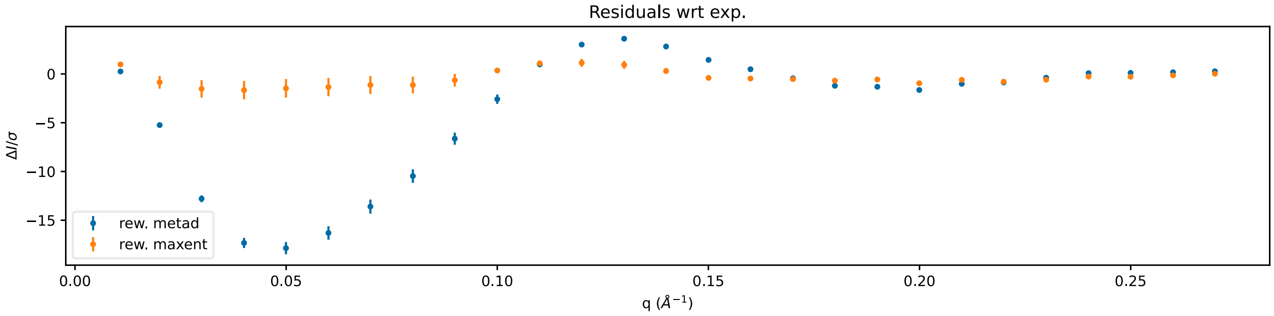


**Figure S11** Residual between computed (metad and maxent-reweighted ensembles, in blue and orange, respectively) and experimental SAXS spectra.

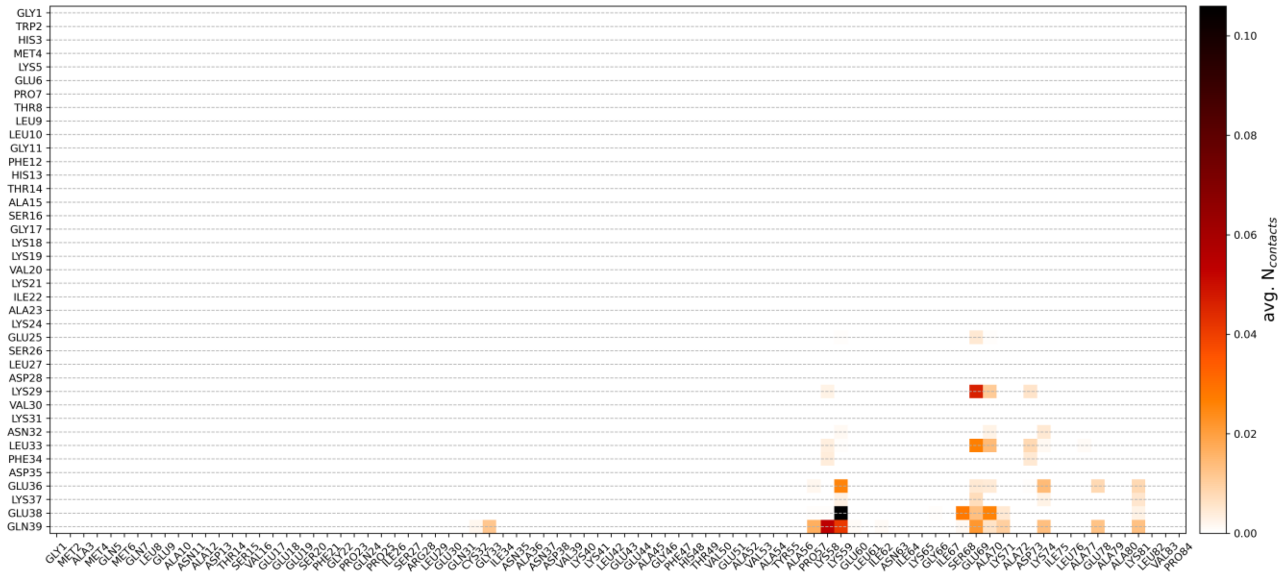

**Figure S12** Residue-wise contact matrix between RAD51's N-ter and BRC4.
